# Pleiotropy and repeated mutation drove convergent domestication of grain amaranth

**DOI:** 10.64898/2026.02.26.708151

**Authors:** Tom S. Winkler, Maxime Kadner, Corbinian Graf, Roswitha Lentz, Julio Martinez, Akanksha Singh, Markus G Stetter

**Affiliations:** Institute for Plant Sciences, University of Cologne, Cologne, Germany; Cluster of Excellence on Plant Sciences, University of Cologne, Cologne, Germany

**Keywords:** Domestication, repeated evolution, seed color, dormancy, pleiotropy

## Abstract

Humans reshaped environments for domesticated plants, yet the extent of human agency in the domestication process remains unresolved. Color variation has been used as evidence for an intended process by humans, as the trait has been mainly linked to visual properties. In the ancient pseudocereal grain amaranth major domestication traits have not changed, but the seed color changed repeatedly from dark to white during multiple domestication processes. Here, we show that the transition to white seed color was of major ecological significance beyond human preference through mechanistic links to seed germination. We identify major metabolomic and transcriptomic changes in the proanthocyanidin pathway leading to loss of seed color and dormancy in domesticated amaranth. Despite the potentially large mutational target size of the proanthocyanidin pathway, the repeated mutation in a single gene led to the domestication phenotype. Multiple independent knock-out mutations in this gene reveal the repeated selection for the loss of seed dormancy in different parts of the Americas. Competition experiments show that the pleiotropic link between seed color and seed dormancy provides a competitive advantage to white seeds in agricultural environments. The reduction of proanthocyanidin content in the seed is observed in numerous crops, including rice and beans, suggesting a common ecological adaptation after early humans altered their environment through changing lifestyles. Our findings establish a direct mechanistic link between individual mutations, pathway-level biochemical reconfiguration, and ecological fitness, illustrating how plants adapted to human-modified environments largely without intentional human agency.

---

The domestication of plants and animals allowed humans to have a sedentary lifestyle and the development of modern societies (1; 2). Their close association with human history and food security underline the importance of the domestication process until today. In addition, domesticated plants and their evolution have long served as example of repeated phenotypic change (3; 4). The cause for the domestication of plants has long been debated by anthropologists, archaeologists, and evolutionary biologists alike (5; 4). While some argue for a conscious and intended selection of specific plants by humans (6) other scholars argue for an evolutionary process where humans only altered environments to which plants adapted (4; 7; 5; 8). Within the last 10,000 years, human impact on environments and organisms around them has strongly increased through the alteration of landscapes and the close interaction with specific animal and plant species (9; 10; 11). At the same time, humans might have intentionally picked plants for specific traits. Revealing the drivers of plant domestication can help to understand human history and improve crops in changing environments. Plant domestication resulted in a suit of convergent phenotypic changes across many crop species, including loss of seed shattering, strong increases in seed size, and frequently seed color changes from darker to lighter seed colors (12; 13; 14). Inferences into the molecular basis of crop domestication have revealed complex examples of convergent and non-convergent genetic changes underlying different adaptive phenotypes in various crop species (15; 16; 14), with contributions from newly arisen mutations and standing genetic variation (13; 17). While many trait changes can be explained by evolution without human intention, some traits have been associated with human preference and conscious decisions because no immediate fitness advantage could be derived from them. Particularly color variations that are observed across species have served as evidence for intentional human selection during domestication (4). Yet, these traits might have other, ecologically relevant functions that increase the fitness of crops in human-made environments.

Grain amaranth is an ancient pseudocereal from the Americas with a high nutritional quality due to the favorable amino acid composition and the gluten free nature of its seeds (18; 19). The grain amaranths were domesticated at least three times independently from their wild relative *Amaranthus hybridus* L., giving rise to the Central American *A. hypochondriacus* L. and *A. cruentus* L., and the South American *A. caudatus* L. with contribution of the endemic *A. quitensis* Kunth. (Fig. 1a; 20; 21). Despite their repeated domestication, some key domestication traits are only weakly pronounced in domesticated grain amaranth. For instance, seed size increased only slightly and crop varieties display strong seed shattering, traits that are altered in all major grain crops (20). However, one trait that was repeatedly altered during amaranth domestication was seed color. The seed coat changed from dark to white in accessions of all three domesticates (21). The selection pressure and molecular basis for the repeated change in this visual trait remains unknown.

**Figure 1.**
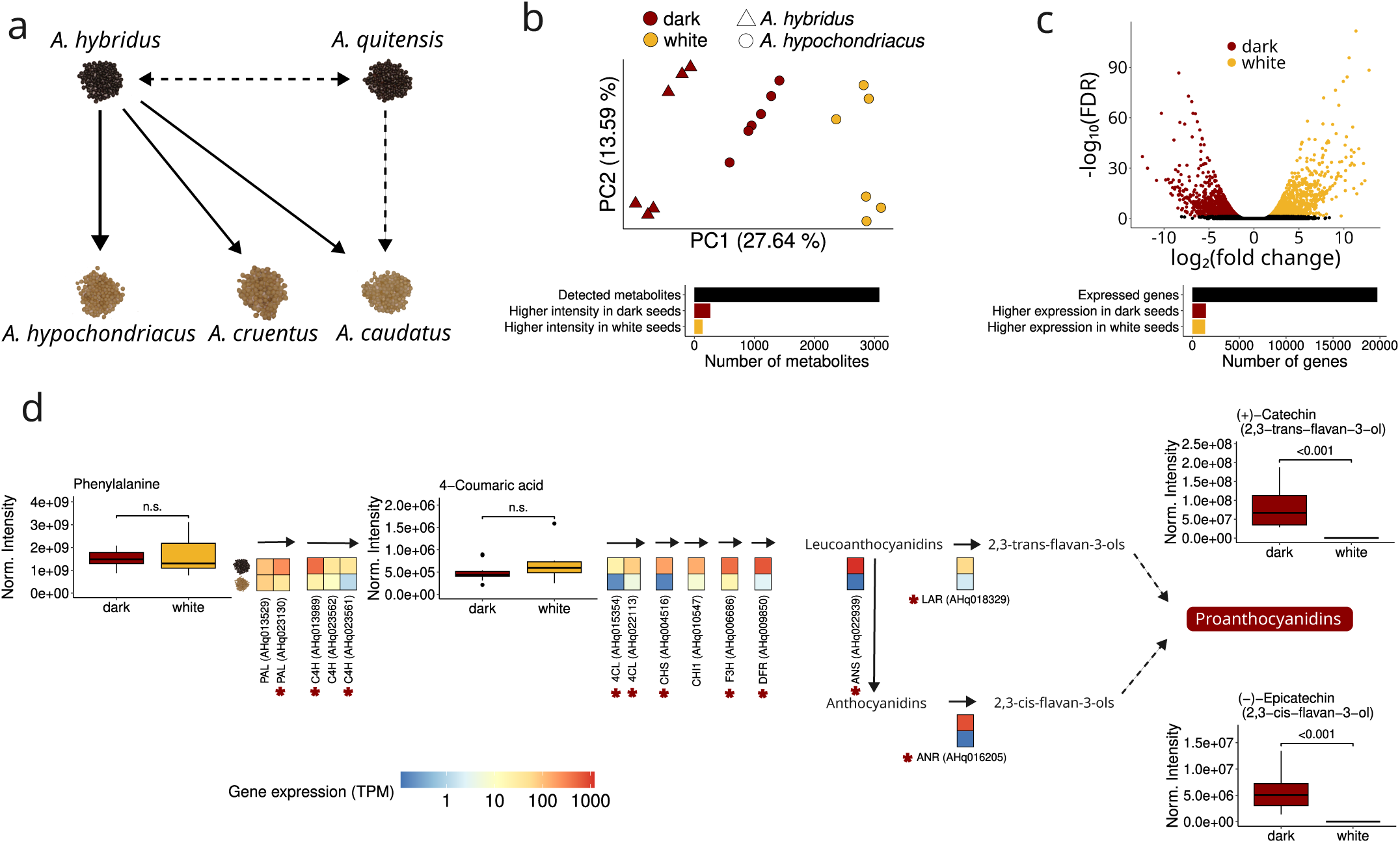
Gene expression differences cause reduced proanthocyanidin accumulation in white seeded *A. hypochondriacus* accessions. **a**: Seed color changed from dark to white during the repeated grain amaranth domestication. The three domesticated species (*A. hypochondriacus*, *A. cruentus* and *A. caudatus*) were domesticated from *A. hybridus*, with contributions from *A. quitensis*. **b**: Principal component analysis using all detected metabolites from dark (dark red) and white (yellow) seeds of *A. hybridus* (triangles) and the domesticated *A. hypochondriacus* (points). **c**: Volcano plot of all expressed genes in developing seed of dark and white seeded *A. hypochondriacus* accessions depicts log_2_-fold change between seed color and -log_10_ of differential expression false discovery rate. Genes with significantly increased expression (absolute log_2_-fold change > 1 and false discovery rate < 0.05) in dark or white seeds were colored dark red or yellow, respectively. The boxplots in b and c depict the number of total detected (black) and differentially expressed metabolites and genes in grain amaranth seeds. Metabolites and genes with significantly higher expression in dark or white seeds were colored dark red or yellow, respectively. **d**: Mean gene expression in Transcripts per Million (TPM) and detected metabolites in developing seeds of *A. hypochondriacus* accessions with dark and white seeds along the general phenylpropanoid and flavonoid pathway leading to proanthocyanidin biosynthesis. Detected seed metabolites are displayed as boxplots of normalized intensity along the pathway. Significant differences between metabolites of two white seeded *A. hypochondriacus* accessions (n = 3 for each) and two dark seeded *A. hypochondriacus* and *A. hybridus* accessions (n = 3 for each) were assessed using a linear model and ANOVA at adjusted p < 0.05. n.s.: non significant. Gene expression is displayed as mean from two dark (n = 7 and n = 3) and white seeded accessions (n = 5 and n = 3), respectively. Genes with significantly higher expression in dark seeds at adjusted p < 0.05 and absolute log_2_-fold change > 1 were annotated using asterisks. No depicted genes had higher expression in white seeds.

In this study we investigate the adaptive advantage of the repeated seed color change during do-mestication, by combining multiomics and evolutionary analyses in the orphan crop grain amaranth. We identify how repeated *de novo* mutations in a key regulator gene changed the expression of many pigment genes to alter seed properties. Ecological and physiological experiments show how the change in seed color enabled the adaptation to human-made environments. Beyond amaranth, loss of seed pigmentation has been described for a range of domesticates and has been used as evidence for intentional selection by humans. However, our results suggest that pale seed color emerged as adaptation to the anthropogenic niche.

## Results

### Reduced proanthocyanidin accumulation determines white seed color in A. hypochondriacus

Seed color is fixed for dark seeds in the wild *A. hybridus*, but polymorphic between accessions of *A. hypochondriacus*. To investigate the molecular basis of amaranth seed color, we measured the seed metabolome of mature seeds of two accessions of *A. hybridus* and two dark- and two white-seeded accessions of the domesticated *A. hypochondriacus* using untargeted LC-MS analysis. We detected a total of 2,917 metabolites, of which 254 could be matched to plant metabolite database entries. Clustering of the metabolites clearly grouped samples according to their domestication status and seed color (Fig. S1). The first principal component of all detected metabolites distinguished samples based on color and species (Fig. 1b), revealing distinct metabolite profiles of the different groups. Seeds of domesticated individuals with dark seeds clustered between the wild accessions and the white accessions, which are likely more domesticated. We identified 402 differentially accumulated metabolites between dark and white seeds while accounting for accession and species as covariates (Fig. 1b). Only one KEGG pathway was enriched in the differential metabolites; the ’Flavonoid biosynthesis’ (adj. p-value = 0.029), which includes the 2,3-flavan-ols (+)-Catechin and (-)-Epicatechin. While early flavonoid pathway metabolites showed no difference in abundance, the accumulation of (+)-Catechin and (-)-Epicatechin was strongly reduced in white seeds (Fig. 1d). The polymerisation of (+)-Catechin and (-)-Epicatechin results in the formation of dark brown proanthocyanidins. These pigments accumulate in the seed coat and confer dark seed color in many plant species after oxidation (22). Hence, the dark seed color in amaranth is most likely determined by the accumulation of proanthocyanidins, which have been lost in the process of domestication leading to the white seed color in domesticates.

The genetic basis of traits can help reveal the selection pressures underlying trait variation and co-emergence. To understand what led to the reduced proanthocyanidin concentration and the consequent seed color change, we compared gene expression patterns between dark and white seeded accessions of *A. hypochondriacus* (Fig. S2). We found a large number of genes differentially expressed between seed colors (14 % of expressed genes; 1,434 higher expressed in dark seeds, 1,358 higher expressed in white seeds; Fig. 1c, Fig. S2). The large number of differentially expressed genes indicates divergence in more traits than only seed color, which is in agreement with the difference in domestication progress between the white and dark seeded accessions. Yet, differentially expressed genes showed strong enrichment for genes related to the phenylpropanoid biosynthesis, which includes the production of flavonoids (Fig. S3, table S1). Consequently, we further investigated gene expression patterns along the flavonoid pathway and related expression with metabolite abundance (Fig 1d, table S2). We found a reduced expression of general phenylpropanoid and early flavonoid pathway genes, including important enzymes (*PAL*, *C4H*, *4CL*, and *CHS*) and of all late flavonoid pathway genes (*DFR*, *LAR*, *ANS* and *ANR*) in white seeds (Figs. 1d, S4). Our metabolomic and transcriptomic analyses show that the reduced accumulation of proanthocyanidins results from a downregulation of the late flavonoid pathway. This change in the expression of nine early and all late flavonoid genes led to the loss of dark seed color during the domestication of amaranth.

### Knockout of the seed color regulator AmMYBL1 leads to white seed color in A. hypochondriacus

The metabolomic and transcriptomic analyses show a highly polygenic signal for the seed color adaptation. Yet, previous work identified few QTL for seed color across five species of grain amaranth and wild relatives (21). Population structure between species potentially lowered the statistical power of the analysis. Thus, we conducted genome wide association mapping with whole-genome sequencing data of 144 *A. hypochondriacus* accessions. We also identified the highest association on chromosome 9, including the previously suggested candidate gene *AmMYBL1* (AHq015545; Fig. 2a; 21). Yet, *AmMYBL1* was not differentially expressed between dark and white seeds (Fig. 2b). However, the phylogenetic position of *AmMYBL1* clusters the protein as R2R3 MYB transcription factor in subgroup S5 of the MYB transcription factors (23). Given the role of subgroup S5 MYB transcription factors in the regulation of proanthocyanidin biosynthesis in other species (24; 25), *AmMYBL1* potentially has a prominent role in seed pigmentation in amaranth.

**Figure 2.**
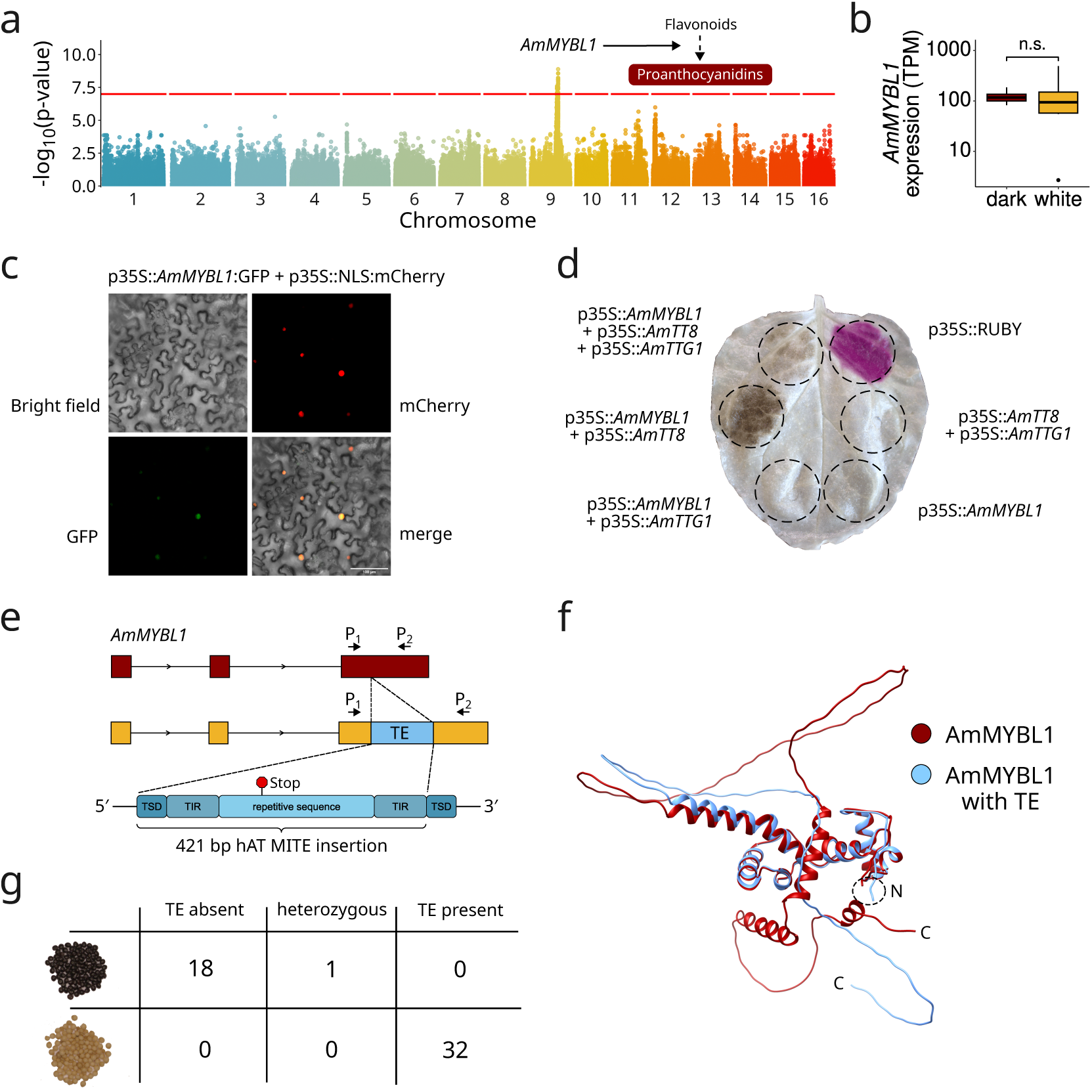
Transposable element insertion into the proanthocyanidin regulator *AmMYBL1* leads to loss of seed color in *A. hypochondriacus*. **a**: Seed color GWAS using 144 *A. hypochondriacus* accessions. The position of *AmMYBL1* on the peak on chromosome 9 was annotated in the plot along its proposed function in proanthocyanidin regulation. **b**: Comparison of *AmMYBL1* gene expression from developing seeds of dark and white seeded *A. hypochondriacus* accessions in Transcripts per Million (TPM). n.s.: non significant. **c**: Subcellular localization of GFP-tagged AmMYBL1 and mCherry fused to a nuclear localization signal in epidermal tobacco cells. Top left: bright field, top right: mCherry channel, bottom left: GFP channel, bottom right: merge of all three channels. Scale: 100 µm **d**: Transient overexpression of *AmMYBL1* and interaction partners leads to dark pigment production in tobacco leaves. Clockwise, from the top right: 1: RUBY positive control, 2: *AmTT8* + *AmTTG1*, 3: *AmMYBL1*, 4: *AmMYBL1* + *AmTTG1*, 5: *AmMYBL1* + *AmTT8*, 6: *AmMYBL1* + *AmTT8* + *AmTTG1*. All genes under the control of CaMV 35S promoter. **e**: Insertion of a 421 bp miniature inverted-repeat transposable element (MITE) of the *hAT* superfamily into the third exon of the *AmMYBL1* gene. Binding location of the primers used for TE genotyping are displayed as P_1_ and P_2_. The premature stop encoded by the TE sequence is annotated. Note that TSD and TIR were magnified for illustration purposes. TSD: target site duplication, TIR: terminal inverted repeat. **f**: Protein structure prediction of *AmMYBL1* without TE (red) and including the inserted TE sequence until the premature stop (blue). The adjacent N-termini and both C-termini were annotated. **g**: TE genotypes from PCR-based genotyping of the *AmMYBL1* TE-insertion for 51 *A. hypochondriacus* accessions of dark or white seed colors.

The mechanistic link between *AmMYBL1* function and pigment accumulation in amaranth seeds was previously unknown. We cloned the *AmMYBL1* CDS from an accession with dark seeds and expressed it in *Nicotiana benthamiana* leaves. The localization of the MYBL1 protein in the nucleus indicates function as expression regulator (Fig. 2c). Nevertheless, the overexpression of *AmMYBL1* alone did not lead to pigment accumulation in *Nicotiana benthamiana* leaves (Fig. 2d). The joint sequence analysis of *AmMYBL1* with subgroup S5 MYBs from other species confirmed the presence of the subgroup-specific sequence-motif (DExWRLxxT; 26) C-terminal of the DNA-binding domains and a conserved bHLH-interaction motif in the R3 domain ([DE]Lx2[RK]x3Lx6Lx3R, Figs. S5 and S6; 27). This conserved motif in *AmMYBL1* and the requirement of proanthocyanidin MYBs in other species to establishment the MBW (MYB-bHLH-WDR) protein complex for transcriptional regulation suggest similar protein-protein interactions in amaranth (27). Therefore, we also cloned the amaranth orthologs of *TT8* (bHLH; AHq013577; Fig. S7) and *TTG1* (WDR; AHq012650; Fig. S8). Yeast two-hybrid assays confirmed that the amaranth MBW proteins interact and can still interact with their *A. thaliana* orthologs (Fig. S9). While neither of the genes alone was sufficient to produce the pigment, co-infiltration of *AmMYBL1* with *AmTT8* and co-infiltration of the complete MBW complex led to an accumulation of dark pigments in *N. benthamiana* (Fig. 2d). In grain amaranth, *AmTTG1* was expressed in most assessed tissues and the highest expression of *AmTT8* and *AmTTG1* was in developing seeds (Fig. S10). The expression of *AmMYBL1* was specific to young and mature seeds (Fig. S10), supporting the control of the flavonoid pathway and the accumulation of proanthocyanidins by *AmMYBL1* together with its interaction partners of the MBW complex.

In addition to the molecular validation of the *AmMYBL1* transcription factor, the identification of the causal mutation for the loss of dark color can help understand the evolutionary history of the trait. Since the gene was not differentially expressed, we investigated the pangenome of grain amaranth (28) to find potential sequence differences between white and dark versions of *AmMYBL1* in *A. hypochondriacus*. The reference genome from a *A. hypochondriacus* accession, which produces white seeds, has a 421 bp insertion of a Miniature Inverted-repeat Transposable Element (MITE) of the *hAT* superfamily in the third *AmMYBL1* exon (Fig. 2e) compared to all non-*A. hypochondriacus* genome assemblies (Fig. S11). The transcription of the inserted transposable Element (TE) would lead to missense mutations and a premature stop resulting in truncation of the last 123 amino acids, representing a strong candidate for loss-of-function of *AmMYBL1* (Fig. 2e). While the truncation still enables correct folding of the N-terminal DNA-binding domains in predicted protein structures, the inserted TE abolishes three predicted alpha helices at the C-terminal part of the protein (Figs. 2f, S12). We investigated whether a truncated AmMYBL1 gene could still induce pigment formation. We generated different AmMYBL1 proteins carrying C-terminal truncations (Fig. S13a,b). The truncated AmMYBL1 proteins were unable to induce pigment formation, confirming that the TE insertion resulted in a functional knockout of the gene (Fig. S13c). In contrast, an in-frame deletion at the site of the TE still enabled pigment accumulation when overexpressed (Fig. S13c). These results show that the C-terminal truncation of *AmMYBL1* through the TE insertion leads to reduced pigment accumulation.

To assess the prevalence of the TE insertion across *A. hypochondriacus* accessions, we performed PCR-based genotyping of the TE in 51 diverse *A. hypochondriacus* accessions. Across accessions, we identified a perfect association between the homozygous TE allele and white seed color (18 dark accessions had no TE, 32 white accessions had the homozygous TE allele; Fig. 2g; table S3). The protein truncation of the color regulator AmMYBL1 through the inserted TE and the association with white seed color in domesticated accessions confirms the TE as causal mutation leading to white seed color in *A. hypochondriacus*. Collectively, our transcriptomic, metabolomic, and molecular analyses show that the insertion of a TE led to the knockout of the positive color regulator *AmMYBL1*, which leads to a downregulation of seed flavonoid genes and reduced accumulation of seed proanthocyanidins, resulting in white seed color of modern *A. hypochondriacus* cultivars.

### Repeated knockout of AmMYBL1 leads to color loss during amaranth domestication

While we find the TE insertion as the causal mutation for seed color loss in *A. hypochondriacus*, we did not find the *AmMYBL1* TE insertion in the other two domesticated amaranth species *A. caudatus* and *A. cruentus*, for which the white seed color is also present (Fig. 1a). The independent domestication of the three crop species could have proceeded through different causal genes or independent mutations in the same gene. If seed color was controlled by different genes, crosses between white-seeded accession of different species would complement the dominant dark seed color. Yet, we did not observe a complementation of the *AmMYBL1* gene in experimental crosses between all three grain amaranth species (table S4), showing that the *AmMYBL1* gene is causal in all white seeded accessions, but through different causal mutations. Hence, the convergent emergence of white seed color in these three domesticates has not occurred through different genes and not through the single *de novo* insertion of the TE that spread through gene flow between domesticates, as the TE is only present in white *A. hypochondriacus* accessions.

Yet, the repeated selection for color-loss alleles already segregating in populations of the wild ancestor or multiple independent *de novo* mutations could have caused the phenotypic change. To investigate the evolutionary history of the seed color change in the grain amaranth species, we generated haplotype networks of the *AmMYBL1* gene sequence from 108 accessions of the wild and domesticated species (Fig. 3a, Fig. S14). While all white seeded *A. hypochondriacus* accessions shared a single *AmMYBL1* haplotype that included the TE insertion, we found the insertion to occur on a network edge specific for *A. hypochondriacus* (Fig. 3a).

**Figure 3.**
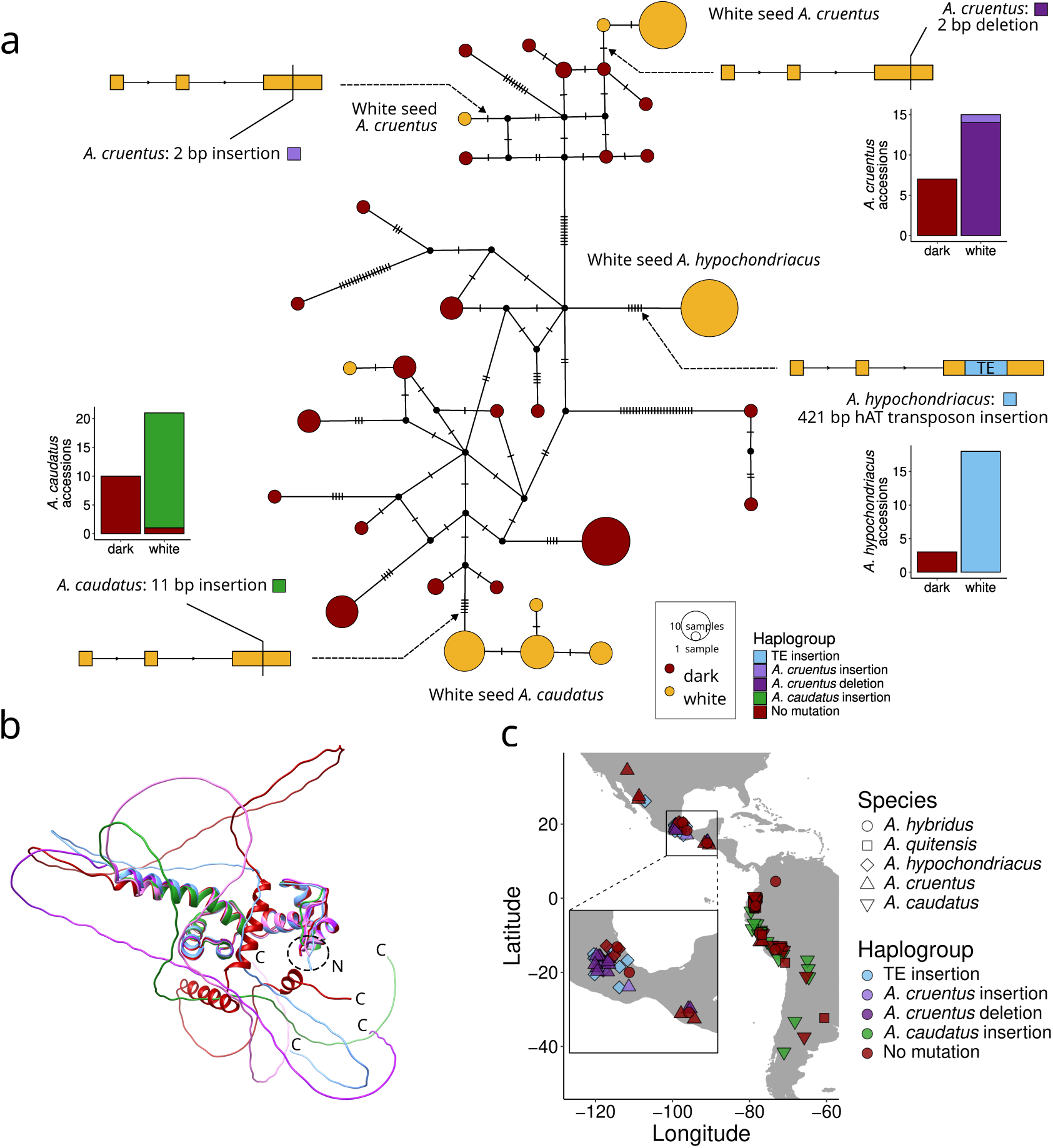
Independent functional mutations in *AmMYBL1* cause white seed color in the three domesticated grain amaranth species. **a**: *AmMYBL1* haplotype network created from 216 haplotypes of the three domesticates (*A. hypochondriacus*, *A. cruentus* and *A. caudatus*) and the two wild relatives *A. hybridus* and *A. quitensis*. The size of the nodes shows the number of identical haplotypes. Mutations are indicated by marks on the edges. The occurrence of high-impact mutations in *AmMYBL1* was annotated on the network and haplotypes were colored according to seed color of the accession. The boxplots show the presence of homozygous functional mutations in *AmMYBL1* in differently colored accessions of *A. hypochondriacus*, *A. cruentus*, and *A. caudatus*. No high-impact mutations were identified in wild accessions. **b**: Protein structure prediction of *AmMYBL1* without functional mutations (darkred) and including all detected functional mutations. The adjacent N-termini and all C-termini were annotated. **c**: Geographic location of 106 accessions with available collection sites. Symbol shape corresponds to the assigned species and symbols were colored according to the haplogroup indicating the presence of homozygous functional mutations in *AmMYBL1*.

We evaluated all accessions for high-impact variants that could impair the function of *AmMYBl1* and lead to color loss. We identified three other segregating high-impact variants in the third exon leading to frameshifts and protein truncations, which were specific to white seeded accessions of the domesticates (Fig. 3a,b, Fig. S12). In the domesticated species *A.caudatus*, we identified an eleven base pair insertion shared by nearly all white seeded *A. caudatus* accessions (20 out of 21 accessions), supporting an independent loss-of-function of *AmMYBL1* in the South American species (Fig. 3a-c). An additional white seeded *A. caudatus* accessions carried an *AmMYBL1* haplotype closely related to dark seeded *A. caudatus* accessions and might represent another loss of seed coat pigmentation. In *A. cruentus*, we identified two distinct high-impact *AmMYBL1* variants associated with white seed color; a two base pair insertion and a two base pair deletion. These mutations did not co-occur in the same individuals and appeared at different edges of the network, suggesting two distinct origins of white seed color in *A. cruentus* (Fig. 3a, S14), coherent with the strong population structure within *A. cruentus* (29). Despite their sympatric occurrence, the *AmMYBL1* haplotypes of white seeded accessions of the two Central American species *A. hypochondriacus* and *A. cruentus* showed clear distinction in the network (Fig. 3a). The shared genetic basis of white seed color from *de novo* mutations in the three different species (albeit with different causal alleles) shows the importance of the trait during amaranth domestication.

### White seed color is associated with reduced seed dormancy in domesticated grain amaranth

The importance of color might be attributed to the visual appearance of the seeds, but proanthocyanidins are also involved in important physiological programs during plant development. Mutants of model plants deficient in seed proanthocyanidins often show altered germination properties and are affected in water and gas permeability and hormonal crosstalk (30; 31). While many wild plants display strong seed dormancy, in crop species where the life cycle is managed by human, fast and uniform germination after sowing might translate into a competitive advantage. White seed color in crops could be associated with faster germination through the loss of proanthocyanidins in the seed coat. To investigate a possible pleiotropic connection between seed color and germination while controlling for the genetic background of the plants, we generated a segregating F_2_ population from a cross between two inbred *A. hypochondriacus* accessions with contrasting seed colors. The seed color of F_2_ lines (n = 511) could be classified into dark (n = 385), brown (n = 87), and white (n = 39) color categories. To assess differences in germination behavior, we recorded seed germination of differently colored F_2_ lines after varying periods of after-ripening. We found significantly higher rates of germination in white and brown seeded lines compared to dark lines from freshly harvested seeds (n = 150, p-value < 2e-16; Fig. 4c). Despite seeds of all colors being viable, dark seeds did not germinate in the tested range, suggesting their higher dormancy. We repeated the germination experiment after four months of storage and all seed colors had higher germination rates than fresh seeds, however dark lines still germinated significantly less than non-dark lines (n = 130, p-value < 2e-16; Fig. 4c). During after-ripening, seed dormancy declined until lines did not differ in germination. After two years of storage in the cold, all seed colors showed similar, uniformly high rates of germination (n = 45, p-value = 0.159; Fig. 4c). The uniformly high rates of germination for all seeds after two years show the high viability of the seeds in the experiment. Our results suggest that the proanthocyanidin accumulation in dark amaranth seeds is associated with significantly higher initial dormancy, while white seeds can germinate even immediately after harvest.

**Figure 4.**
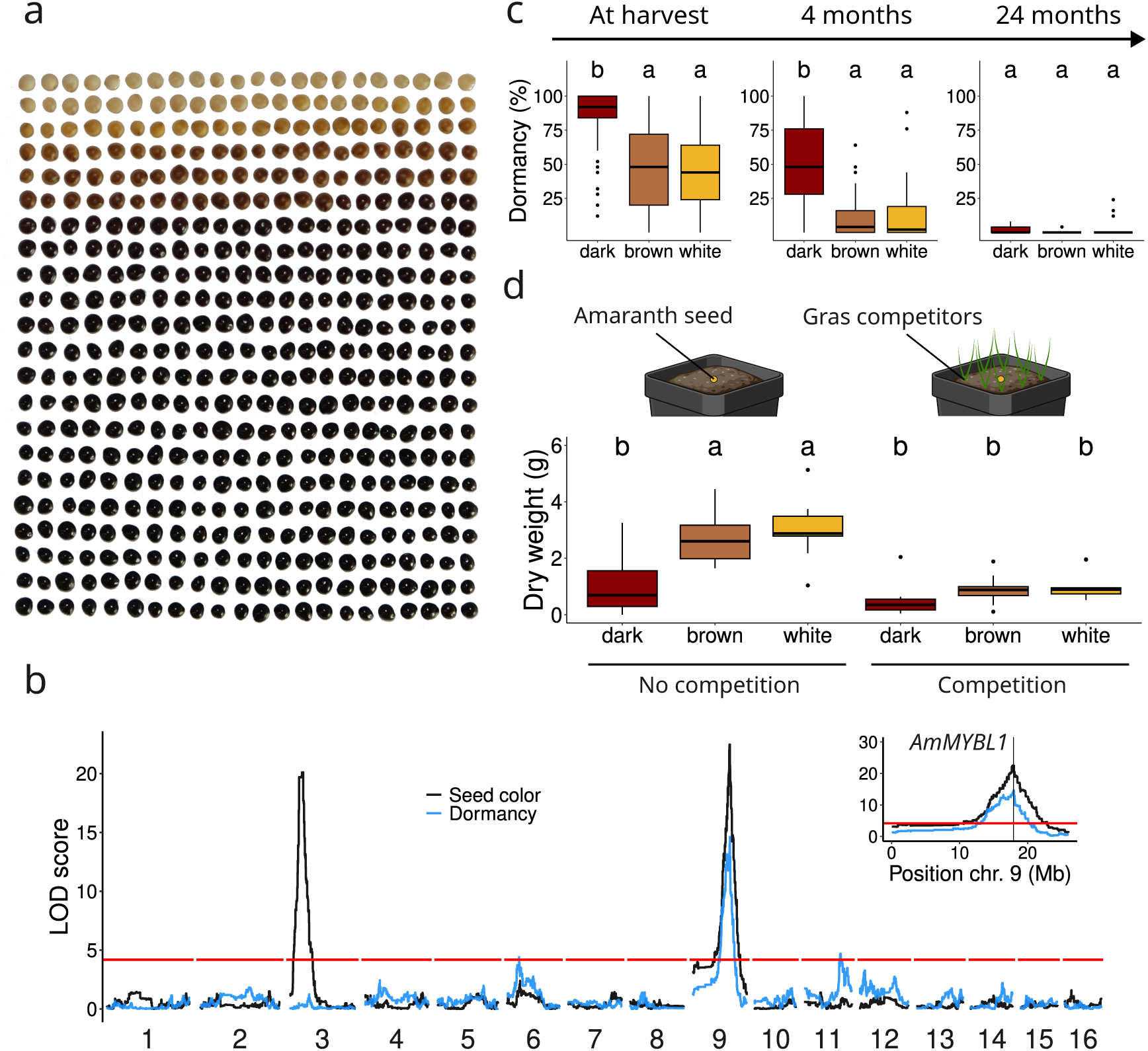
Pleiotropic control of seed color and germination behavior in *A. hypochondriacus*. **a**: Seed color diversity in *A. hypochondriacus* F_2_ mapping population. Each seed represents one F_2_ line. **b**: QTL mapping for results for seed color (black) and germination rate (blue) in *A. hypochondriacus* F_2_ population along the 16 chromosomes. The horizontal red line depicts LOD significance threshold for QTL based on 1000 permutations. Insert depicts closeup of chromosome 9, with the position of the seed color regulator *AmMYBL1* marked by a vertical line. **c**: Germination rate of white, brown and dark seeded F_2_ lines. Germination experiments were conducted immediately after harvest (n = 150), after four months (n = 130) of storage and two years of storage (n = 45). Significant differences between seed colors were assessed using ANOVA and Tukey’s test at p < 0.05. **d**: Dry weight of white, brown and dark seeded F_3_ families (10 families per seed color, 10 replicates per family and condition) in presence or absence of inter-species competition seven weeks after sowing. Significant differences between seed colors were assessed using ANOVA and Tukey’s test at p < 0.05.

To investigate the genetic control of seed properties in our mapping population, we used linkage mapping in the bi-parental population. The segregation ratio of seed color fit a two-locus inheritance model with dominant epistasis (Fig 4a; F_2_ phenotypic segregation ratio 12:3:1, Chi-square p-value = 0.303, table S5). The dominant dark color allele at the one locus can ’mask’ the phenotype at a second hypostatic locus, where a dominant brown allele controls the seed color only if the first epistatic locus is in the recessive state. The dominant epistasis was further supported by phenotypic segregation of selfed F_2_ lines in the F_3_ generation, where dark and brown seeded F_2_ lines could change to produce white seeds, while already white seeded lines (inferred to be homozygous for the recessive allele at both causal loci) exclusively produced white seeds (Fig. S15). While seed color could be classified into discrete categories, seed germination followed a quantitative trait distribution (Fig. S16). To identify the underlying loci controlling dormancy, we produced whole-genome sequencing data from 493 lines from the F_2_ population. This dense genotyping allowed us to identify a total of 230,125 biallelic SNPs between the two parental lines (Fig. S17). The mapping of seed dormancy identified QTL on chromosome 6 and 11 and a major QTL on Chr. 9 (one LOD support interval from maximum likelihood estimate, chr. 9: 17.791-17.971 Mb) strongly overlapped the *AmMYBL1* seed color QTL (chr. 9: 17.655-17.971 Mb; Fig. 4b). Seed color in the cross was controlled by two QTL, on amaranth chromosomes 3 (controlling brown color) and 9 (controlling loss of a dark seed coat and including *AmMYBL1* (Fig. 4b, Fig. S18; 21), fitting the previously identified two-locus dominant epistasis inheritance model. The overlap of the QTL for seed color and dormancy further confirms the pleiotropic control of the two traits by *AmMYBL1* in grain amaranth.

Loss of seed dormancy and fast germination could provide a competitive advantage to crops in agricultural environments through improved early vigor. To test whether differences in germination behavior between differently colored lines could translate into performance advantages in an agricultural environment, we planted seeds of differently colored F_3_ families either in absence (to simulate a human managed growth environment) or presence of a grass competitor and recorded plant dry weight seven weeks after sowing as proxy for plant performance. F_3_ families with white and brown seed color showed both, faster germination and reduced dormancy in soil compared to dark seeded families (Fig S19, S20). The improved germination resulted in significantly increased above-ground biomass for non-dark F_3_ families in the absence of competition compared to the same families with competitors (Fig. 4d, S21). Our soil grown experiments show that loss of dark seed color in amaranth likely led to an improved performance in agronomic environments through the reduction in seed dormancy. The ecological advantage of white seeds might explain the repeated selection of knockout alleles of the same regulator gene in the three domesticates.

## Discussion

### Parallel evolution during domestication

The history of crops and humans is tightly linked and the integration of evidence from archaeobotanical, genetic and theoretical studies has shaped our understanding of crop domestication (32). For instance, archaeobotanical studies revealed millenia-long transition periods of wild barley and einkorn wheat morphotypes to the domesticated forms (33) and there is evidence for important roles of polygenic selection from standing genetic variation and gene flow from the wild relative (34; 35). Modeling of the evolutionary trajectories of domestication alleles in adzuki bean, maize and rice suggests that mutations could predate the split between wild ancestor and domesticate (17; 36; 37). In the repeatedly domesticated common bean and grain amaranth, only few common genes were under selection in distinct gene pools, suggesting distinct genetic pathways toward domestication (38; 21). However, the molecular basis of crop evolution led to the repeated identification of convergent genetic changes at orthologous genes in different species underlying key domestication phenotypes (15; 39; 40; 41). Our haplotype reconstruction of the *AmMYBL1* locus supports soft selective sweeps at the seed color QTL and consolidates the independent domestication processes of the three domesticates (Fig. 3; 21). Contrary to other crops, we did not detect any of the causal alleles in the wild relatives, indicating that seed color and dormancy adaptation resulted from repeated *de novo* mutations during the domestication of grain amaranth in different regions of the Americas. While strongly reduced dormancy is likely maladaptive in wild species, color-loss alleles could predate the domestication and be maintained in the wild relative at low frequency in a heterozygous state. This has been found for the wild ancestor of maize, where the open fruit case is likely deleterious, but the maize allele likely predates domestication (36). Yet, the four independent mutations within 149 bp in the same gene that seem to not be present in the wild relative, suggest repeated seed color adaptation from *de novo* mutations. Our results support the view of domestication as complex process, in which adaptation simultaneously acted on new mutations and standing genetic variation leading to molecular convergence or distinct genetic responses between populations depending on selective pressure and trait architecture.

### Pale seed color as target for selection during domestication

Adaptation of key domestication traits has previously been proposed to have resulted from uninten-tional selection during onset of domestication for phenotypes, beneficial under agricultural practices (42; 43). While the reduction of seed dormancy and shattering may increase fitness in agricultural environments directly, other phenotypes may change due to indirect selection or pleiotropic control with selected traits (43; 44). For instance, in vegetable crops, for which the seed is not a directly yield-associated organ, seed size increased comparable to grain crops (43), which may be explained by unintentional selection for the improved germination and seedling vigor of larger seeds during early domestication. In contrast, the loss of dark seed color observed in many crops has been proposed to be due to selection based on cultural preference and taste. The loss of flavonoid-based seed pig-mentation is found in diverse crops, including *Brassica rapa* (45), rice (46), bean (40), adzuki bean (17), and quinoa (47), suggesting its selection during domestication. The co-localization of QTL of seed dormancy and proanthocyanidin accumulation demonstrate the pleiotropic control of the two traits (Fig. 4). Our results indicate that selection against seed dormancy and for improved germination probably led to seed color changes through the pleiotropic control of seed color and germination. Our ecological experiments show the advantage of pale seed color in agricultural environments. Rapid seed color change and improved germination also advanced the *de novo* domestication of pennycress as it allows uniform cultivation (48). Although selection due to cultural preferences could explain individual transitions to pale seed color for certain regions and time periods, it fails to account for the frequent seed color transitions in diverse crop species originally domesticated in different domestication centers among different time frames (12; 4). Instead, repeated selection for improved germination properties could explain convergent seed color changes in diverse crop species. Given the wide-spread transitions to pale seed color in diverse crop species during domestication, selection against the pleiotropic controlled seed dormancy is the likely causal selective pressure on the visual trait.

Our results demonstrate that the reduction of dark colored proanthocyanidins in crop seeds can represent an adaptation to agricultural environments. While the reduction of seed dormancy represents an important adaptation to the anthropogenic niche (9), the overall flavonoid biosynthesis pathway needs to stay intact and generally functional due to pleiotropic constraints in other plant tissues. Based on their selection history and genetic variation, for individual crop species the loss of proanthocyanidins might lead to the development of maladaptive pre-harvest sprouting (49; 50). The potentially large mutational target size for loss of proanthocyanidins in the flavonoid pathway that we observe in the metabolomic and transcriptomic results (Fig. 1), is likely reduced due to severe pleiotropic effects. These pleiotropic constraints could partially explain the repeated selection of causal mutations in a single regulator with tissue-specific expression (*AmMYBL1*; Fig. S10) during amaranth domestication. However, the extremely small range of only 149 bp outside the DNA-binding domain in which we find the knock out mutations is surprising and might indicate additional functions of the transcription factor beyond the control of flavonoid biosynthesis.

The repeated loss of seed color from *de novo* mutations during grain amaranth domestication is an example of convergent adaptation to anthropogenic environments. While seed coloration represents the obvious visual trait, selection probably acted on the loss of seed dormancy, which dragged seed color changes along. The observed seed color changes in many crop species, that have been thought of as an example of agency by humans over plant domestication, likely resulted from the pleiotropic control of seed color and dormancy. Domestication can well have proceeded as adaptation of the crop population to a newly created environment, without requiring an active domesticator.

## Acknowledgments

We thank Vivien Plückthun for her contributions to the germination and competition experiments. Figure 4 was partially created in BioRender. Stetter, M. (2026) https://BioRender.com/0vy74ge.

## Author contributions

MGS conceived the research program and coordinated the analyses. TSW analyzed the genomic, transcriptomic and metabolomic data. MK and JM performed molecular analysis of *AmMYBL1*. CG generated the amaranth mapping population and TSW performed the linkage mapping. RL and AS generated sequencing libraries. AS performed GWAS and PCR genotyping of *AmMYBL1*. TSW performed the germination experiments. TSW and MK prepared the figures. TSW and MGS wrote the draft of the manuscript with input from all authors. All authors read and approved the manuscript.

## Data, code and materials availability

All data is available in the main text or the supplementary materials. Code used for analysis is available at: https://git.nfdi4plants.org/stetter-lab/amaranth_seed_color. All generated sequencing raw data is available under Bioproject accession PRJEB106936.

## Supplementary materials

Materials and Methods

Figs. S1 to S23

Tables S1 to S10

## Materials and Methods

### Metabolic analysis of amaranth seeds

To investigate the metabolic basis of seed color in grain amaranth, we performed metabolic profiling of hydrophyllic metabolites in *A. hybridus* accessions PI 511754 and PI 667158, dark seeded *A. hypochondriacus* accessions PI 604581 and PI 604587, and white seeded *A. hypochondriacus* accessions PI 558499 and PI 643070 with three replicates per accession (table S6). For metabolite extraction, 50 mg of dry seed material were prepared using a modified protocol from Salem et al. (2016) (51). Hydrophyllic analytes were measured with a Waters ACQUITY Reversed Phase Ultra Performance Liquid Chromatography (RP-UPLC) coupled to a Thermo-Fisher Exactive mass spectrometer using C18 columns. Chromatograms were recorded in Full Scan MS mode (Mass Range [100-1500]). All mass spectra were acquired in positive and negative ionization modes. Matching criteria for compound annotation were 5.5 ppm and 0.085 minutes deviation from reference compounds mass-to-charge ratio and retention time, respectively. Extraction and mass spectrometry was performed by metaSysX (Potsdam, Germany). We required metabolites to be detected in at least 4 samples to be considered for the subsequent analyses, leaving 2917 metabolites. For the multivariate analysis, metabolite intensities were log_10_ transformed, centered and scaled. For the univariate analysis, we replaced missing values with half the minimum value per metabolite and identified differential metabolites between seed colors using the following linear mixed-effect model: *log*_10_(*metabolite intensity*) ∼ *Seed color* + *Species* + (1|*Accession*).

### Differential gene expression analysis from developing seeds

We isolated total RNA from developing seeds of the dark seeded *A. hypochondriacus* accessions PI 604581 and PI 604587 (seven and three replicates) and the white seeded accessions PI 558499 and PI 643070 (five and three replicates) using the Invitrogen™ PureLink™ Plant RNA Reagent (Thermo Fisher Scientific, catalog number: 12322-012) following the manufacturer’s protocol. Before nucleic acid extraction, plants were grown under well-watered short-day conditions at 60 % air humidity with night and day temperatures set to 19 ^◦^*C* and 24 ^◦^*C*, respectively. We collected developing seeds from the flower with forceps to avoid sampling of flower tissue and immediately stored samples at -80 ^◦^*C* until RNA extraction. Sequencing libraries were prepared using the TruSeq stranded mRNA preparation kit and samples were sequenced on the Illumina NovaSeq 6000 platform (100 bp reads, paired and stranded) by Novogene (Netherlands). We trimmed adapter sequences from the reads using Trimmomatic v0.39 (52) using the ILLUMINACLIP module with the following settings: custom_adapters.fa:2:30:10 (table S7). We mapped the trimmed reads to the *A. hypochondriacus* reference genome v3.0 (53) using STAR v2.7.8a (54) with default settings (table S8). We performed quality control of reads and mapping using FastQC v0.11.9 (55) and Qualimap v2.2.2a (56). We quantified read pairs per gene using featureCounts v2.0.6 (57), counting only read-pairs for which both reads were mapped to the genome. We performed differential gene expression analysis using the variancePartition R package (58) using the formula ∼ *Seed color* + (1|*Accession*) + (1|*Batch*). Genes with absolute log_2_-fold change > 1 and false discovery rate < 0.05 were considered to be differentially expressed. We conducted gene ontology (GO) term enrichment analysis of differentially expressed genes using the goseq R package (59) with the functional annotation of Graf et al. (2025) (53). We used KIPEs v3.2.4 (60) to annotate the general phenylpropanoid and flavonoid pathway genes in the amaranth genome (23; 53). We quantified gene expression in different tissues using a publicly available dataset (BioProject Accession PRJNA263128; 61) of the white seeded *A. hypochondriacus* accession PI 558499 as described in Graf et al. (2025) (53).

*Plant material and sequencing of A. hypochondriacus F_2_ population*

To generate a segregating *A. hypochondriacus* mapping population, we crossed the dark seeded accession PI 604581 and the white seeded accession PI 558499. 493 plants of the F_2_ population were grown in greenhouse under short-day conditions at 60 % air humidity with night and day temperatures set to 19 ^◦^*C* and 24 ^◦^*C*. We sampled plant material from young leaf tissue and extracted DNA using NucleoSpin Plant II kit (Macherey-Nagel) according to manufacturer’s instructions. We generated sequencing libraries using a modified Nextera shallow-sequencing protocol as described in Singh and Stetter (2025) (62). Libraries from 493 F_2_ individuals and both parental accessions were sequenced on Illumina NexSeq with 2×150 bp by Novogene, United Kingdom.

*Assessment of germination properties and competition experiment from A. hypochondriacus mapping population*

We assessed seed color and germination properties of the seeds produced by the *A. hypochondriacus* mapping population described above. We visually classified the seeds of 507 F_2_ lines into three seed color categories (white, brown and dark). To assess germination properties of the seeds produced by the F_2_ population, we recorded the germination of 25 seeds per line over 10 days using a semi-automated setup (Fig. S22). Seeds were placed on wet filter paper (Ahlstrom-Munksjö, Germany) for germination tests in a climate chamber at 22 ^◦^*C* under constant light condition and 60 % air humidity. We recorded images of seeds every 2 h using a custom script and marked the time of germination for each seed using ImageJ (63). We defined the percent of dormant seeds per line as fraction of seeds which did not germinate during the 10 day experiment and counted germinated seeds when the emerging radicle was clearly visible. We tested germination properties from the same sets of seeds at different timepoints, once immediate after harvest (80 dark lines, 41 brown lines, 29 white lines), after four months of storage (67 dark lines, 37 brown lines and 26 white lines), and after two years of storage (15 lines per seed color). We tested for differences in germination behavior between lines of different seed colors using an ANOVA with the design formula: *Median time until germination or logit*(*Percent dormant seeds*) ∼ *Seed color* + *Batch* + *Seed color* : *Batch*.

For the competition experiment, we picked 10 random F_3_ families per seed color category (dark, brown, or white seed color) which were propagated from the F_2_ population described above. At the start of the experiment, individual amaranth seeds were sown into 12×12 cm pots randomised in a randomized complete block design and grown under well-watered long day conditions either with or without competition, with 10 replicates per family and condition. For plants grown with competition, at the start of the experiment eight 10 day old *Dactylis glomerata* seedlings were transplanted as competitors evenly spaced around the focal amaranth seed (Fig. S23). We recorded dry weight at 7 weeks after sowing as proxy for plant performance and assessed significant differences using a linear mixed-effects model with the design formula: *Dry weight* ∼ *Seed color* + *Competition* + *Seed color* : *Competition* + (1|*Line*) + (1|*Block*).

### Linkage mapping of color and germination

We performed quality control of sequencing data from 493 F_2_ lines and the two resequenced parental accessions using FastQC v0.11.9 (55). Next, we mapped reads to the *A. hypochondriacus* reference genome v3 (53) using bwa-mem2 v2.2.1 (64). We marked duplicates using Picard v2.27.5 (65) and called variants using GATK v4.2.2.0 (66) following ’GATK best practices’, resulting in a total of 545,680 biallelic SNPs before filtering. We filtered the dataset for SNPs with contrasting homozygous calls in both parental lines and allele frequency 0.15-0.85 across all samples using bcftools v1.19 (67), retaining 496,932 SNPs. To correct for random genotyping errors and bias in calling heterozygous genotypes due to low-coverage sequencing, we imputed variant calls through the reconstruction of recombination breakpoints using NOISYmputer v1.0.0 (68). The recombination breakpoint reconstruction led to 230,124 imputed SNPs. We used the imputed genotypes to perform QTL mapping for seed color and germination behavior phenotypes using R/qtl2 (69), defining QTL based on 95 % LOD significance thresholds from 1000 permutations.

### Genome-wide Association Analysis and TE genotyping

To assess the species-specific association of seed color, we performed GWAS with 144 classified accessions of *A. hypochondriacus* from around the globe described in Stetter et al. (2020) (21) and Sing and Stetter (2025) (62). We processed the raw reads and called variation as described previously, retaining only biallelic SNPs and discarded variants with more than 30% missing data. We further imputed the missing genotypes using Beagle v5.5 (70) and filtered genotypes with minor allele frequency <0.05 resulting in 582,215 SNPs. We performed GWAS in GAPIT (71) using the CMLM model. Both kinship matrix and the first three principal components were used to control for the population structure.

To check the association of TE with the seed color we genotyped 53 diverse accessions of *A. hypochondriacus* from Singh and Stetter (2025) (62). We conducted PCR amplification of the third exon of the *AmMYBL1* gene using primers flanking the 421 bp TE insertion region (forward primer - *AAGCGATTGAACGCAACCAA*, reverse primer *GCTCTGATATTAGCTTAAAATCCCCC*). For PCR amplification, we used the KAPPA2G Robust PCR kit with GC buffer (cat No. KK5004) with program cycle as: 95°C for 5 min, 30 cycles of 95°C for 40 s, 60°C for 30 s, 72°C for 1 min and final extension at 72°C for 5 min. The genotype was accessed using length polymorphism of the amplicons (379 bp without TE and 800 bp with TE) on 2% agarose gel.

### Convergence analysis and haplotype network

To check for seed color complementation in interspecific amaranth crosses, we evaluated the seed color produced by F_1_ plants described in Gonçalves-Dias et al. (2023), (table S4; 72). For the haplotype network analysis, we used public whole-genome sequencing data from 115 native range accessions described previously (table S9; 21). We processed reads and called variants using Picard v2.27.5 and GATK v4.2.2.0 as described previously. For sample-level quality control, we retained only biallelic SNPs and further removed variants with missing data >= 0.3 and minor allele frequency < 5 % using vcftools v0.1.16 (73) resulting in 5,551,849 SNPs. To visualize genetic structure in the dataset, we performed LD pruning using the setting *–indep-pairwise 50 10 0.5* retaining 809,562 SNPs and calculated PCA using plink 1.9 (74). Taxonomic assignment of amaranth accessions based on morphology could lead to erroneous population assignments. To accurately classify the amaranth accessions into their respective species, we reassigned species labels according to their clustering in the genome-wide PCA (table S9, Fig. S24).

The *A. hypochondriacus* reference genome contained a 421 bp TE-insertion in the seed color regulator *AmMYBL1* that could lead to bias in read mapping and variant calling for accessions without this structural variant. To exclude this bias in variant calling, we generated an altered reference genome for which we manually removed the inserted TE sequence and recalled variants. We extracted all mapped reads within 1 Mb up- and downstream of the *AmMYBL1* gene for 108 reclassified accessions with high coverage and available seed color phenotypes from the previously described dataset using Bazam v1.0.1 (75). We remapped all extracted reads against the TE-less *A. hypochondriacus* reference genome using bwa-mem2 v2.2.1 and called variants as described previously. To call larger structural variants missed by GATK, we used Delly v1.2.6 (76) to call and genotype structural variants in the remapped reads. To increase genotyping accuracy, we manually added the 421 bp *AmMYBL1* TE insertion we removed from the reference genome to the Delly database before genotyping. We merged precise structural variants identified with Delly with SNP and indel calls from GATK using bcftools concat. We phased and imputed missing data in this joint dataset using beagle 5.4 (70) and annotated variant effects on protein coding genes using SnpEff v5.1 (77). To construct the *AmMYBL1* haplotype network, we extracted *AmMYBL1* fasta sequences (exons and introns) for 216 haplotypes using vcf2fasta (https://github.com/santiagosnchez/vcf2fasta). Before haplotype network construction, we filtered singleton haplotypes leaving 205 haplotypes. We constructed median-joining networks of *AmMYBL1* haplotypes using PopART (78) with *ɛ* = 0. We predicted 3D protein structures for AmMYBL1 with functional mutations using the server-based implementation of Alphafold3 (79, accessed on 06.08.2025) using AmMYBL1 protein sequences from the following accessions: PI 558499 (*A. hypochondriacus* with and without TE), PI 511680 (*A. caudatus*), PI 576481 (*A. cruentus* insertion), and PI 649623 (*A. cruentus* deletion). We aligned and visualized the predicted protein structures using UCSF chimera v1.19 (80).

### Molecular validation of AmMYBL1

To confirm *AmMYBL1* as potential MYB proanthocyanidin regulator of subgroup S5, we aligned proanthocyanidin MYBs from different species (table S10) with amaranth, *Beta vulgaris*, and *A. thaliana* orthologs of subgroup S5 and S6 (81; 82; 23) using Clustal Omega v1.2.4 (83) and constructed a phylogenetic tree using the BioNJ algorithm implemented in SeaView v5.1 (84) with 1000 bootstrap replicates. To identify amaranth orthologs of MBW complex interactors, we used *A. thaliana* bHLH proteins of subgroup IIIf (85) and the *A. thaliana* WDR protein TTG1 in blast searches against annotated amaranth proteins using blastp v2.12.0 (86) with an E-value cutoff of 1e-10. We aligned amaranth candidates together with *A. thaliana* proteins using Clustal Omega v1.2.4 (83) and constructed neighbour-joining phylogenetic trees with 1000 bootstrap replicates using ClustalW v2.1 (87). Generated trees were visualized using SeaView v5.1 (84).

MBW-complex genes of amaranth (AHq015545, *AmMYBL1*; AHq012650, *AmTTG1*; AHq013577, *AmTT8*) were cloned from cDNA of developing seeds of dark seeded accession PI 604581. Full-length CDS was synthesized using the RevertAid™ H Minus First Strand cDNA Synthesis kit (Thermo Fisher Scientific, cat. no. K1631) according to the manufacturer’s protocol. They were subsequently cloned into plant expression vectors pAMPAT including a CaMV35S promoter sequence using the Gateway cloning system (Thermo Fisher Scientific. (2003) Gateway® Technology. Waltham, MA). The RUBY reporter (88) was cloned into a Greengate expression vector following the standard protocol (Addgene, Kit #1000000036) (89). All constructs were confirmed using whole plasmid sequencing (ONT Lite Whole Plasmid Sequencing, Eurofins). Transient expression assays in *Nicotiana benthamiana* were conducted using a modified protocol by Jörgens et al. (2010) (90). Briefly, we transformed *Agrobacterium tumefaciens* strains GV3101 pMP90RK (for Gateway-constructed vectors) and GV3101 pMP90 (for Greengate-constructed vectors) using the heat-shock method with the binary plasmid pSOUP. *Nicotiana benthamiana* leaves were then transiently transformed using these *A. tumefaciens* strains co-infiltrated 1:1 (v/v) with the antisilencing strain RK19 (91), using a syringe without needle, into the abaxial side of the leaves of 3-4 week-old tobacco plants. Pigment accumulation was documented seven days after infiltration using a Canon EOS 2000D DSLR. Regarding the subcellular localization assays, an expression plasmid containing amaranth AmMYBL1 with C-terminal GFP was assembled into a Greengate expression vector following the standard protocol (Addgene, Kit #1000000036) (89). As positive control for subcellular localization, we used a 35S:SV40NLS:3xmCherry construct in GreenGate plant expression vector (23). Three days post infiltration, confocal laser scanning microscopy was performed on Leica DM5500 with the DCS-SPE (ACS APO 20.0×0.60 water-immersion objective). GFP and mCherry were excited at 395 nm and 587 nm and emission was detected at 509 nm and 610 nm, respectively. To avoid cross talk between excited fluorophores, we performed sequential scanning between frames.

To test their interactions, MBW interactors (AHq015545, *AmMYBL1*; AHq012650, *AmTTG1*; AHq013577, *AmTT8*) were cloned into Gateway™ compatible pAS2-1 and pACT2 yeast expression vectors contain-ing the GAL4-binding domain and GAL4-activator domain respectively (92). Four to six replicates of transformed *S. cerevisiae* line AH109 (93) with every combination of MBW interactors were grown overnight in selective medium SD lacking Leucine (Leu) and Tryptophan (Trp). Cultures were then stamped onto selective SD medium lacking Leu, Trp and Histidine (His), supplemented with 0mM, 5mM or 20mM of 3-amino-1, 2, 4-triazole (3-AT). After 5 days at 30°C, a representative replicate per combination was picked and documented.

## Supplement

### Supplementary Figures

**Figure S1.**
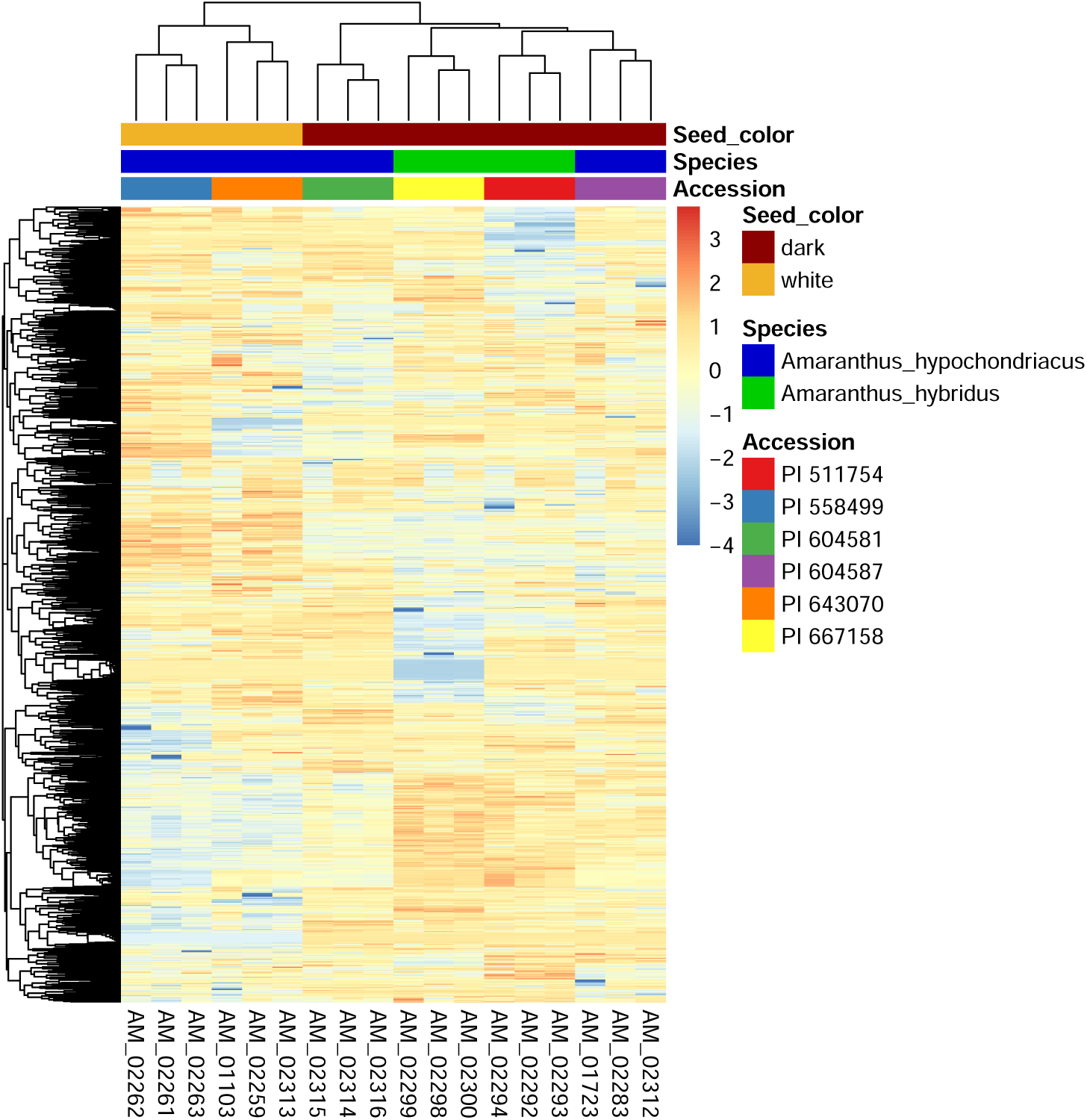
Clustered heatmap of detected metabolites in mature *A. hypochondriacus* and *A. hybridus* seeds. The metabolite heatmap is based on z-scores of the log_10_ transformed normalized metabolite intensity + 1. Indicated on top for different columns are the accession, species and seed color for individual samples.

**Figure S2.**
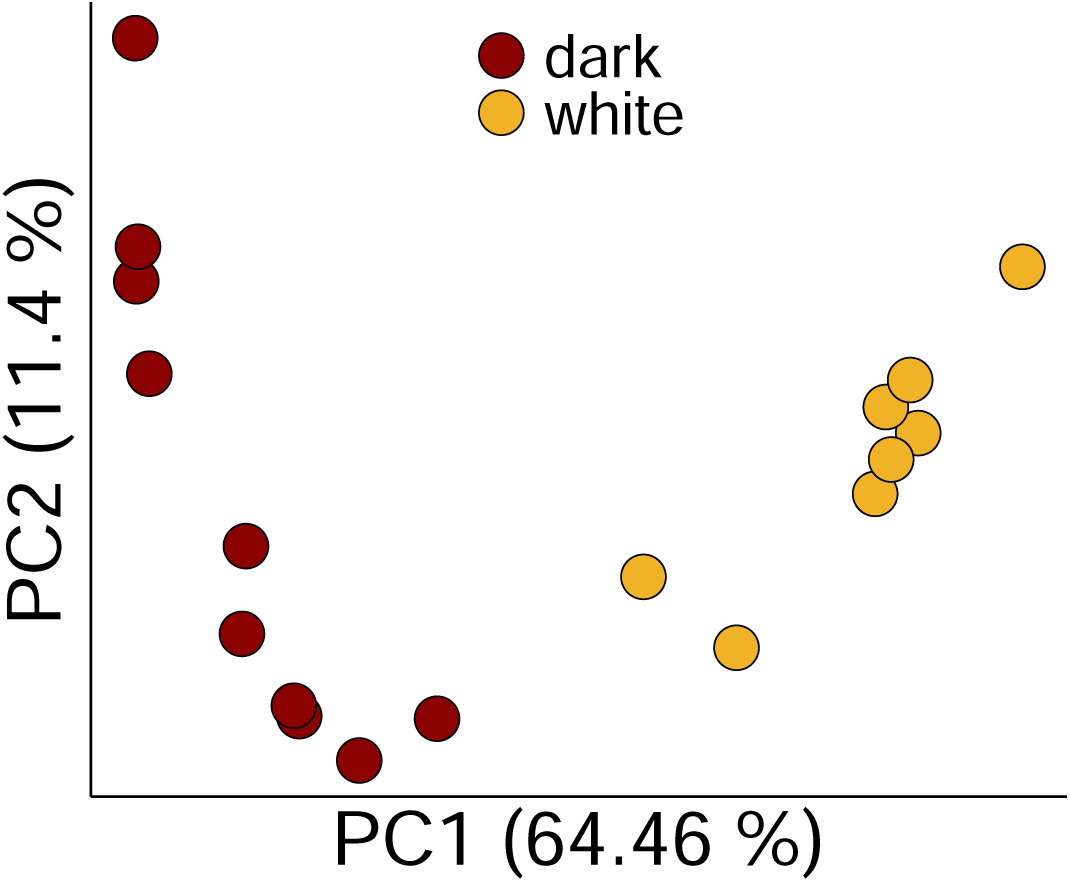
Principal component analysis of the transcriptome of developing seeds of *A. hypochondriacus* accessions with dark or white seeds. PCA was computed from the regularized logarithm transformed normalized counts of the different RNA-seq samples and colored according to the seed color of the accession.

**Figure S3.**
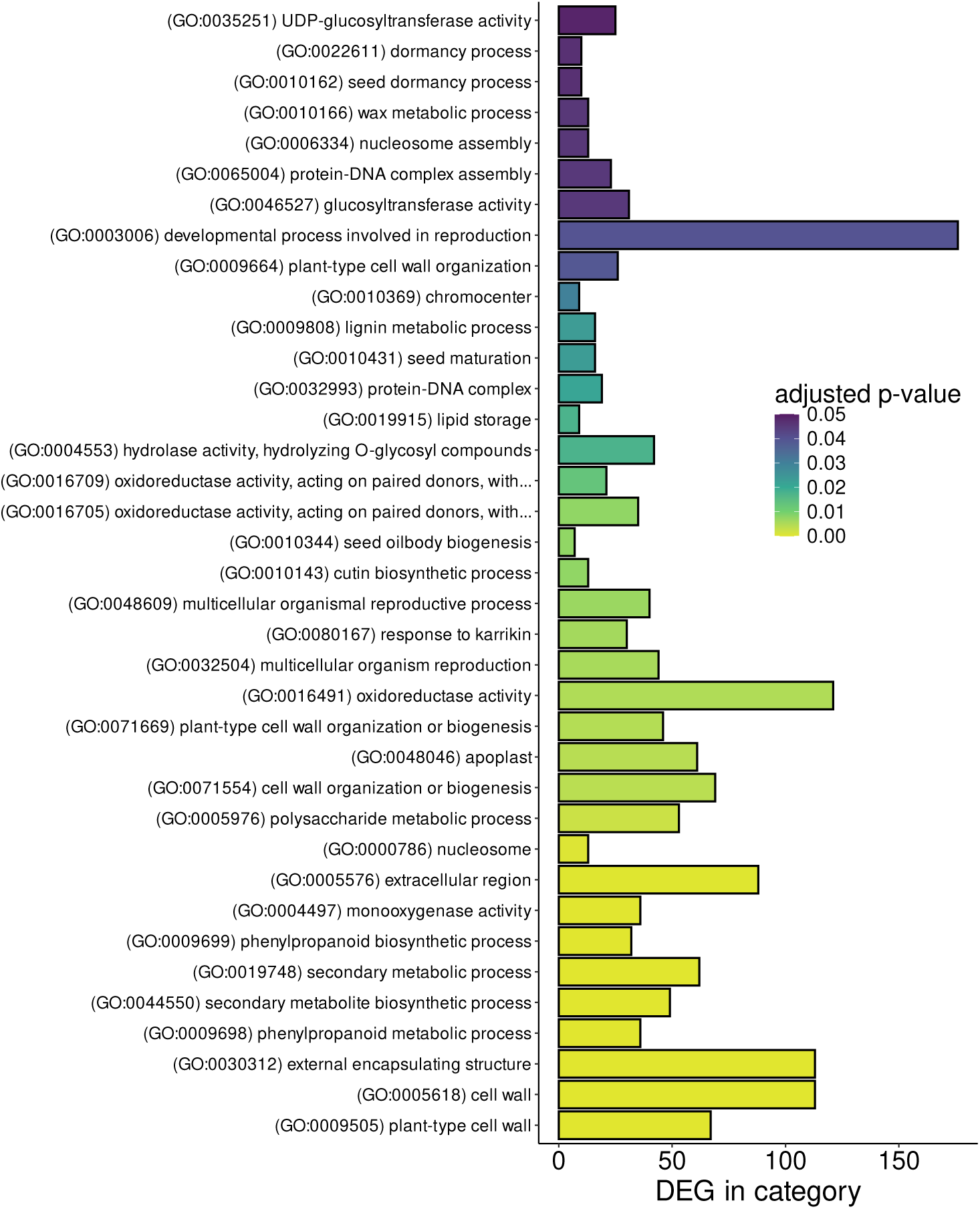
Gene ontology (GO) terms enriched among differentially expressed genes in differently colored *A. hypochondriacus* seeds. Enrichment of GO-terms was calculated comparing all differentially expressed genes against all expressed genes. The x-axis shows the number differentially expressed genes (DEG) per GO category.

**Figure S4.**
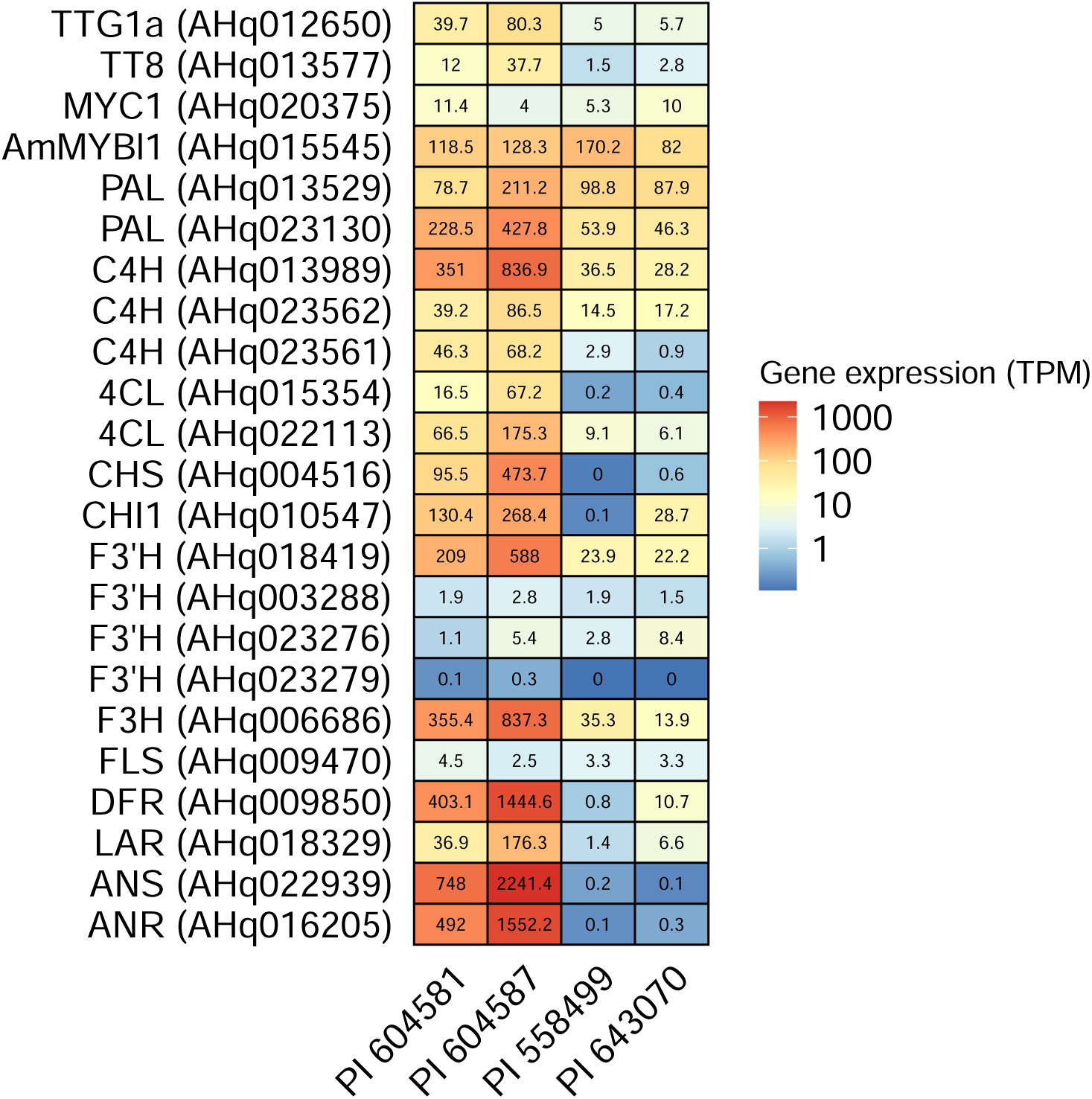
Expression of flavonoid pathway genes in developing seeds of different *A. hypochondriacus* accessions. Expression in Transcripts per Million (TPM) for flavonoid pathway genes in the dark seeded accessions PI 604581 (n = 7) and PI 604587 (n = 3) and the white seeded accessions PI 558499 (n = 5) and PI 643070 (n = 3). Expression values denoted in the tiles are given as mean of all biological replicates for each accession.

**Figure S5.**
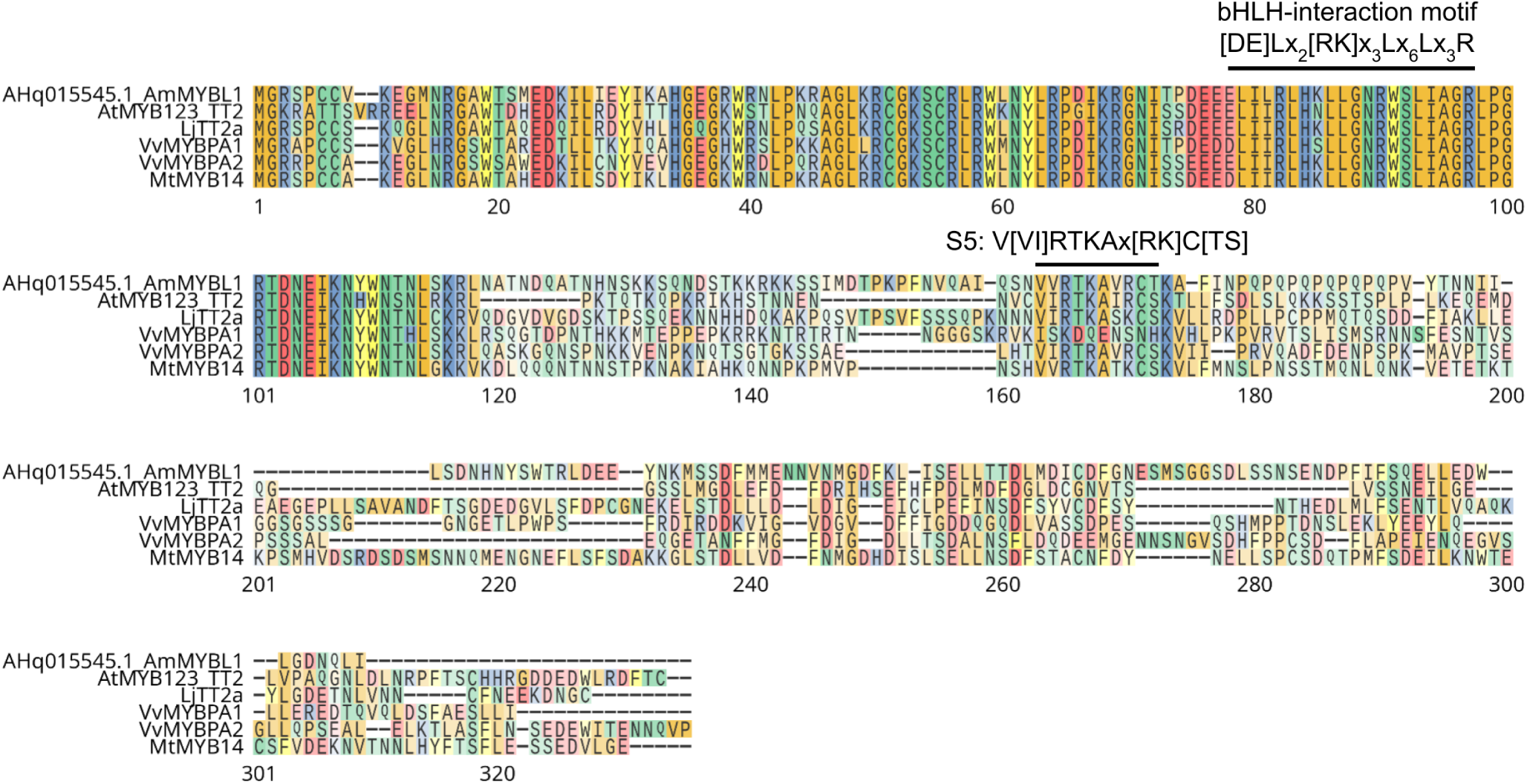
Sequence alignment of S5 MYBs of different plant species. Multiple sequence alignment of S5 proanthocyanidin regulators of different plant species.

**Figure S6.**
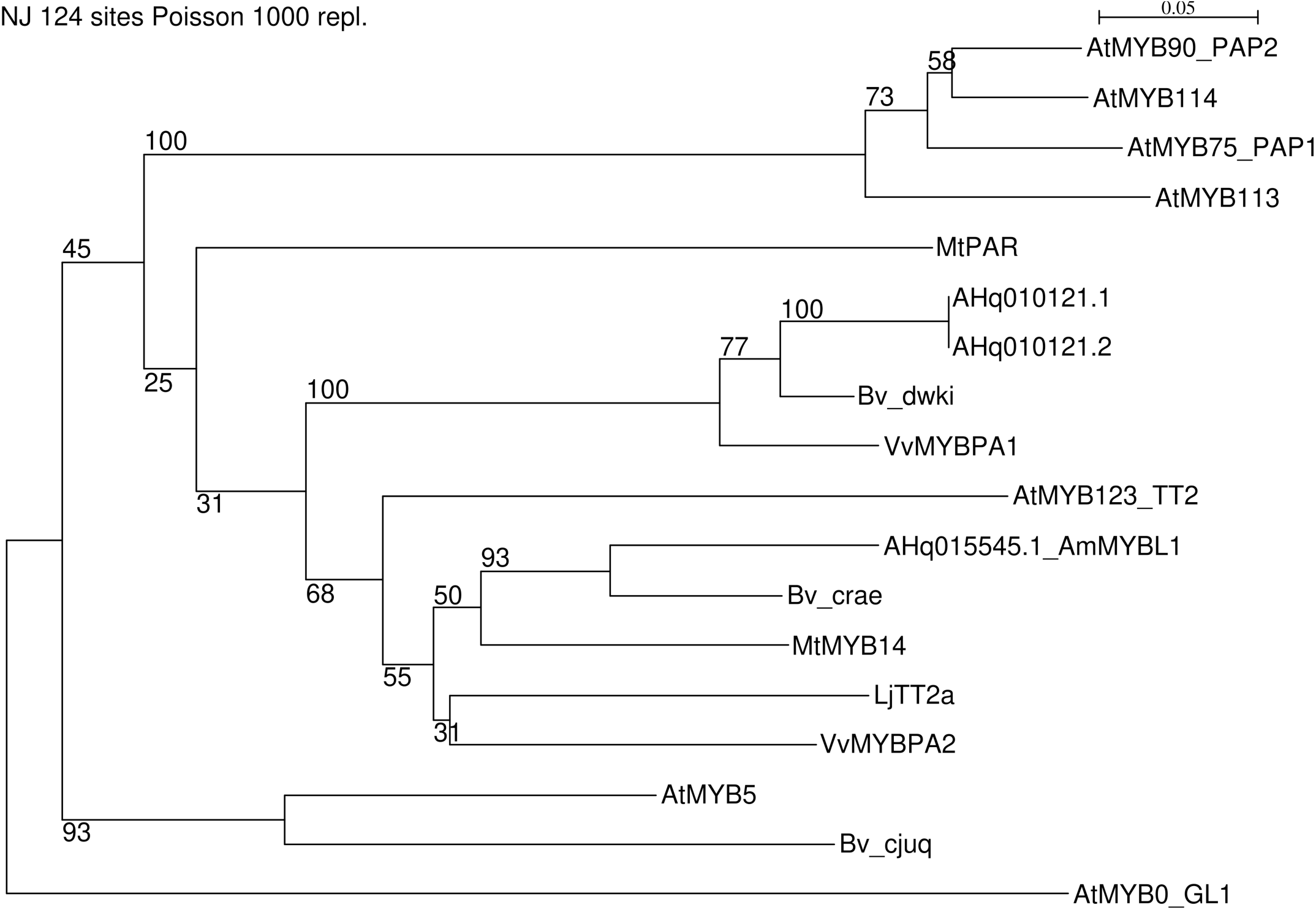
Phylogenetic analysis of *A. hypochondriacus* subgroup S5 MYBs. Neighbour-joining phylogenetic tree of *A. hypochondriacus* MYB proteins of subgroup S5 with *A. thaliana* MYBs of subgroups S5 and S6. S5 MYB proanthocyanidin regulators from other plant species were included in the phylogeny. *A. thaliana* GL1 was used as an outgroup. Branch support from 1000 bootstrap replicates was annotated in the phylogeny.

**Figure S7.**
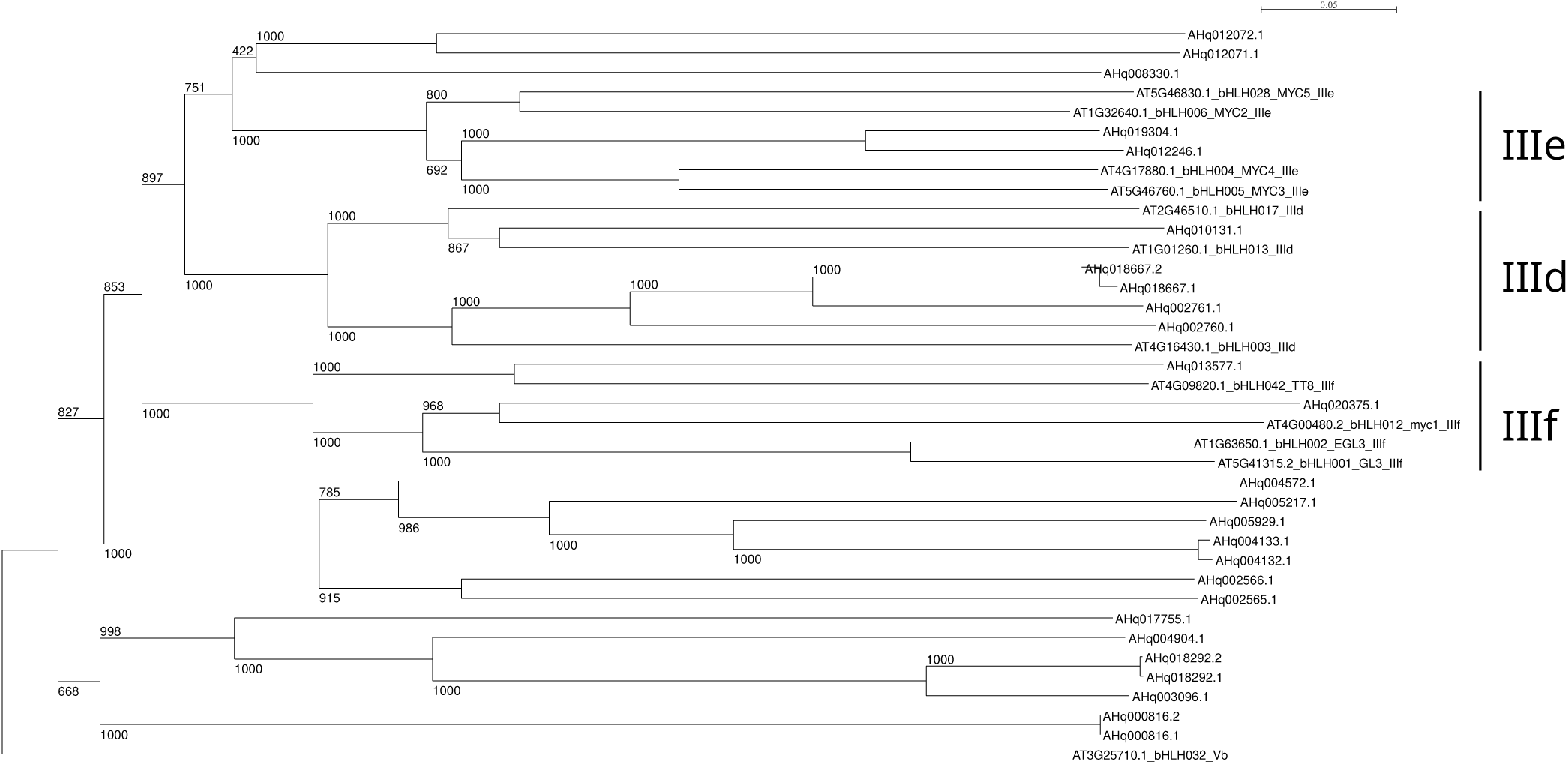
Phylogenetic analysis of *A. hypochondriacus* subgroup III bHLH proteins. Neighbour-joining phylogenetic tree of *A. thaliana* bHLH proteins of subgroups IIId, IIIe and IIIf with ama-ranth candidates. Branch support from 1000 bootstrap replicates was annotated in the phylogeny.

**Figure S8.**
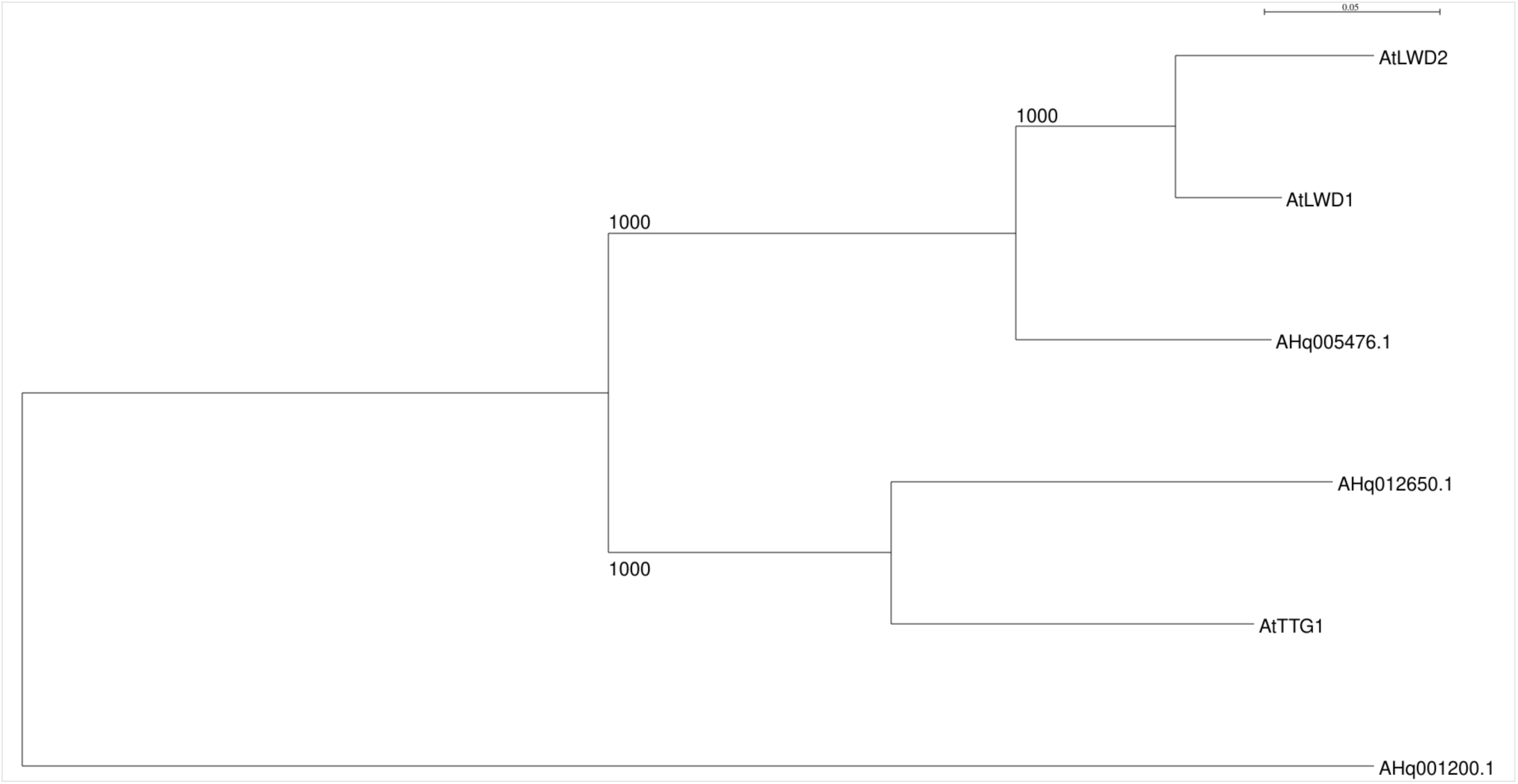
Phylogenetic analysis of *A. hypochondriacus* TTG1 orthologs with *A. thaliana* proteins. Neighbour-joining phylogenetic tree of *A. thaliana* TTG1 and closely related WDR proteins with amaranth candidates. Branch support from 1000 bootstrap replicates was annotated in the phylogeny.

**Figure S9.**
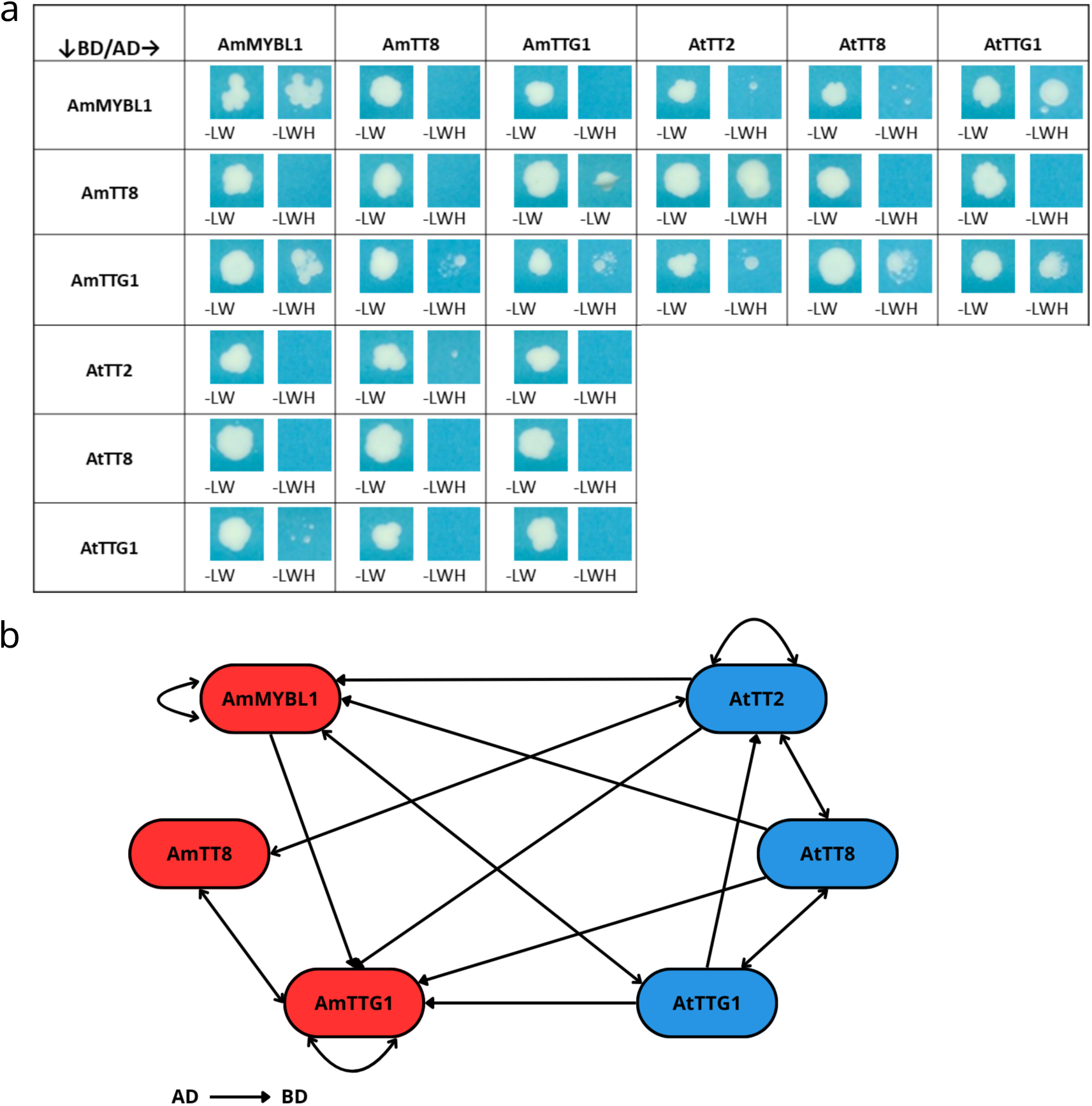
Protein-protein interactions between MBW complex proteins of amaranth and *Ara-bidopsis*. **a**: Yeast two-hybrid assay showing protein-protein interactions between *A. hypochondriacus* (Am) and *A. thaliana* (At) MBW complex proteins. Interaction is indicated by growth of yeast colonies on selective medium SD -LWH, with SD -LW as positive control. Proteins of interest were either fused to the binding domain (BD, rows) or activation domain (AD, columns) to test for pairwise interactions. Yeasts were grown for 5 days. **b**: Summary of detected protein-protein interactions in yeast two-hybrid assays. Single-headed arrow show uni-directional interaction. Double-headed arrows show bi-directional interaction. Interaction data for *A.thaliana* was derived from Baudry et al. (2005) (24).

**Figure S10.**
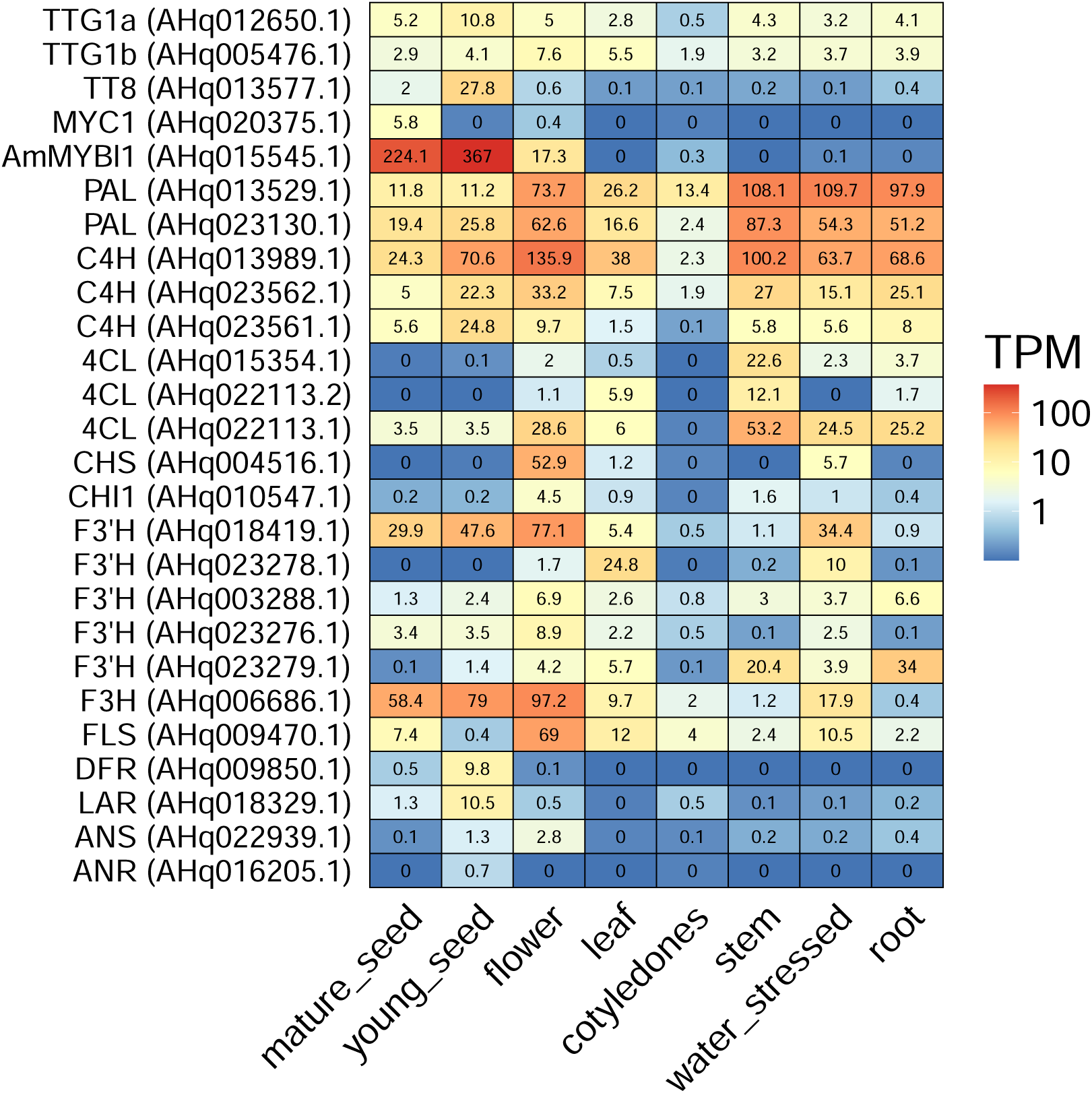
Gene expression of flavonoid pathway genes and regulators in different tissues of *A. hypochondriacus* reference accession PI 558499. Normalized gene expression in TPM of different flavonoid pathway genes and regulators in different tissues and under water-stressed conditions.

**Figure S11.**
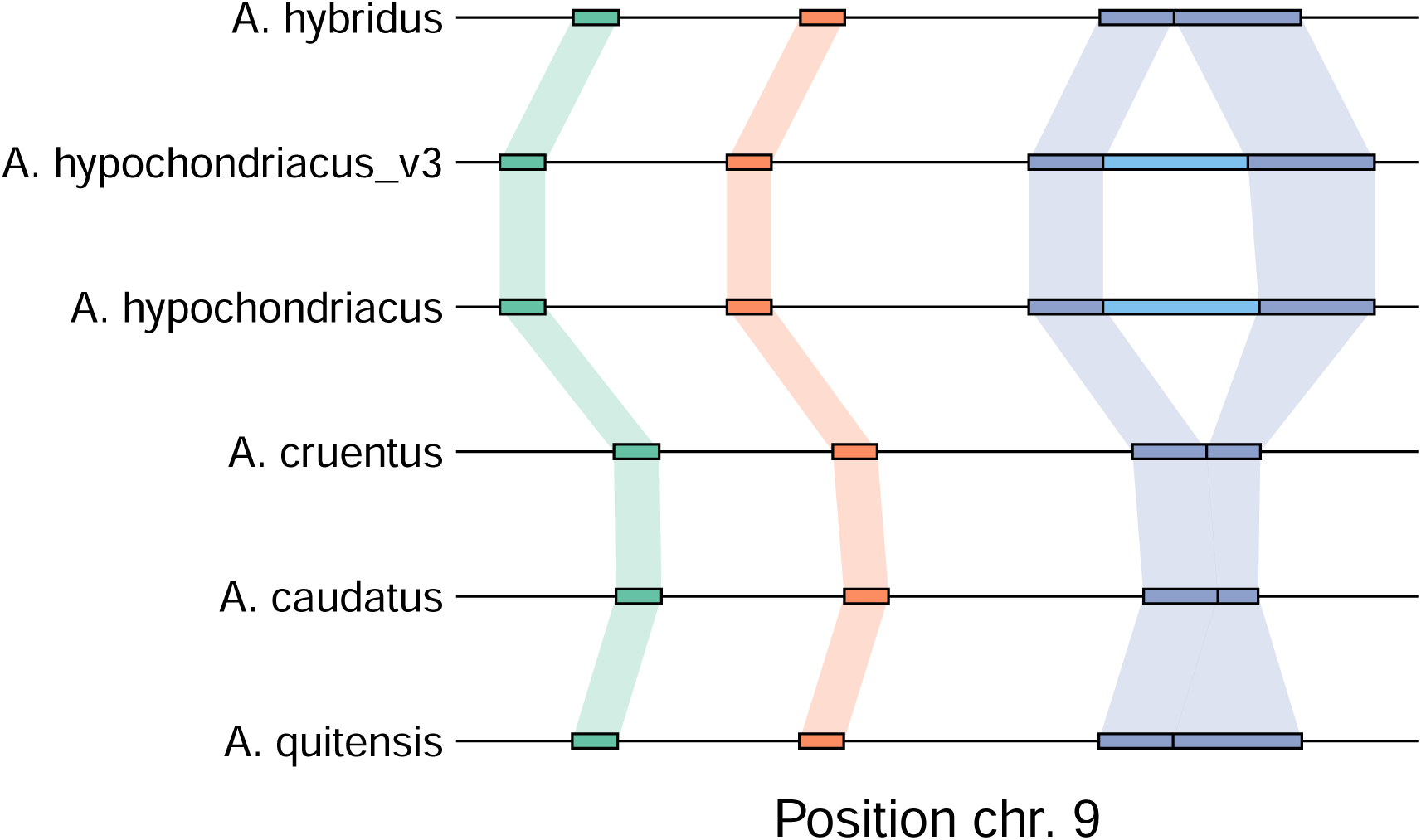
Local synteny of the *AmMYBL1* locus between assemblies of the grain amaranth pangenome. Syntenic relationship between the three conserved *AmMYBL1* exons (green, orange and dark blue) in the different genome assemblies. The inserted transposable element (light blue) is found exclusively in the genomes of white seeded *A. hypochondriacus* accessions. The assemblies correspond to the following accessions: *A. hybridus*, PI 511754; *A. hypochondriacus* v3, reference genome described in Graf et al. (2025) (53), PI 558499; *A. hypochondriacus*, PI 643036; *A. cruentus*, PI 643058; *A. caudatus*, PI 642741; *A. quitensis*, PI 490466.

**Figure S12.**
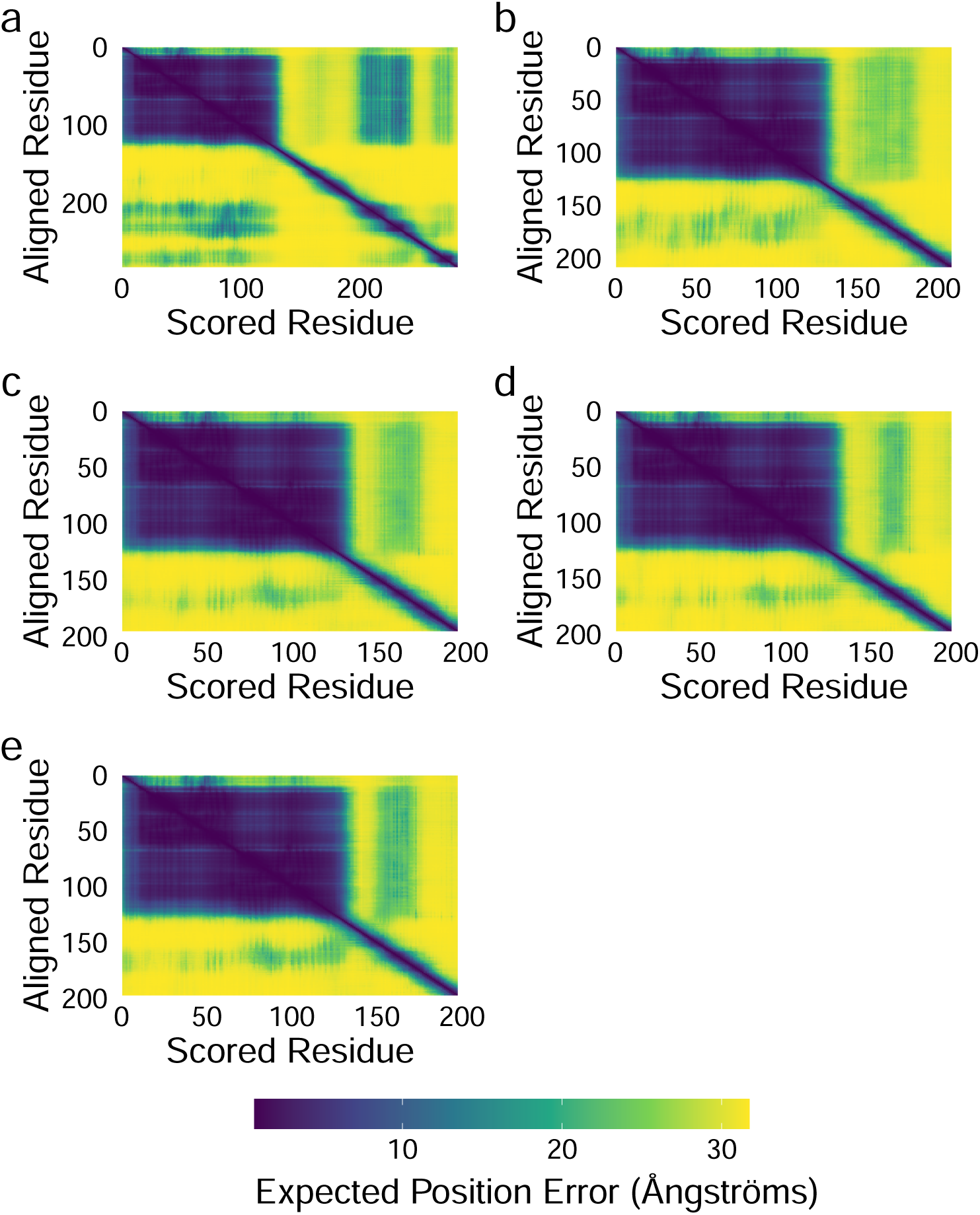
Predicted alignment error for AmMYBL1 protein structure predictions. **a**: Expected position error for aligned and scored residues of AmMYBL1 protein structure prediction. **b**: Expected position error for aligned and scored residues of AmMYBL1 protein structure prediction with inserted TE. **c**: Expected position error for aligned and scored residues of AmMYBL1 protein structure prediction with *A. caudatus* insertion. **d**: Expected position error for aligned and scored residues of AmMYBL1 protein structure prediction with *A. cruentus* insertion. **e**: Expected position error for aligned and scored residues of AmMYBL1 protein structure prediction with *A. cruentus* deletion.

**Figure S13.**
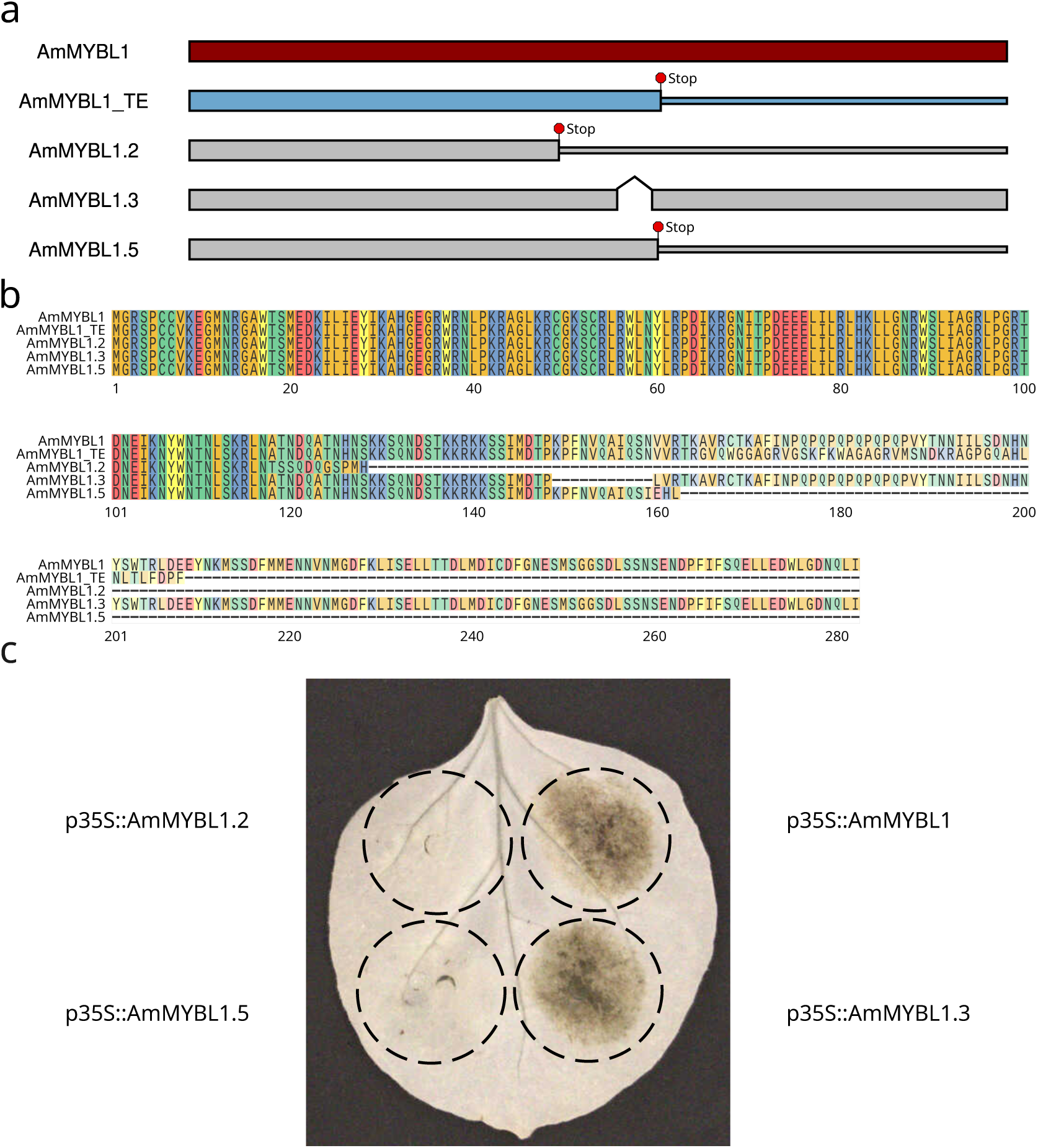
Functional analysis of mutated AmMYBL1 proteins. **a**: Display of CDS structure of functional AmMYBL1 protein from dark seeds, AmMYBL1 with inserted TE and the three mutated AmMYBL1 proteins. Full boxes represent the ORF. Position of stop codons leading to C-terminal truncation are displayed. The in-frame 33bp deletion is displayed as a connector line. **b**: Multiple sequence alignment of AmMYBL1 protein sequences from PI 558499 without inserted TE (AmMYBL1), from PI 558499 with translated TE (AmMYBL1_TE) and with the three mutated AmMYBL1 proteins. **c**: Transient overexpression of truncated AmMYBL1 proteins in *N. benthamiana* leaf via *Agrobacterium* infiltration. Each infiltration additionally included the AmMYBL1 bHLH and WDR interaction partners under the control of CaMV 35S promoter (p35S:AmTT8 and p35S:AmTTG1). Clockwise, from the top right: 1: p35S:AmMYBL1 as positive control, 2: p35S:AmMYBL1.3 (11 AA in frame deletion), 3: p35S:AmMYBL1.5 (C-terminal truncation), 4: p35S:AmMYBL1.2 (C-terminal truncation).

**Figure S14.**
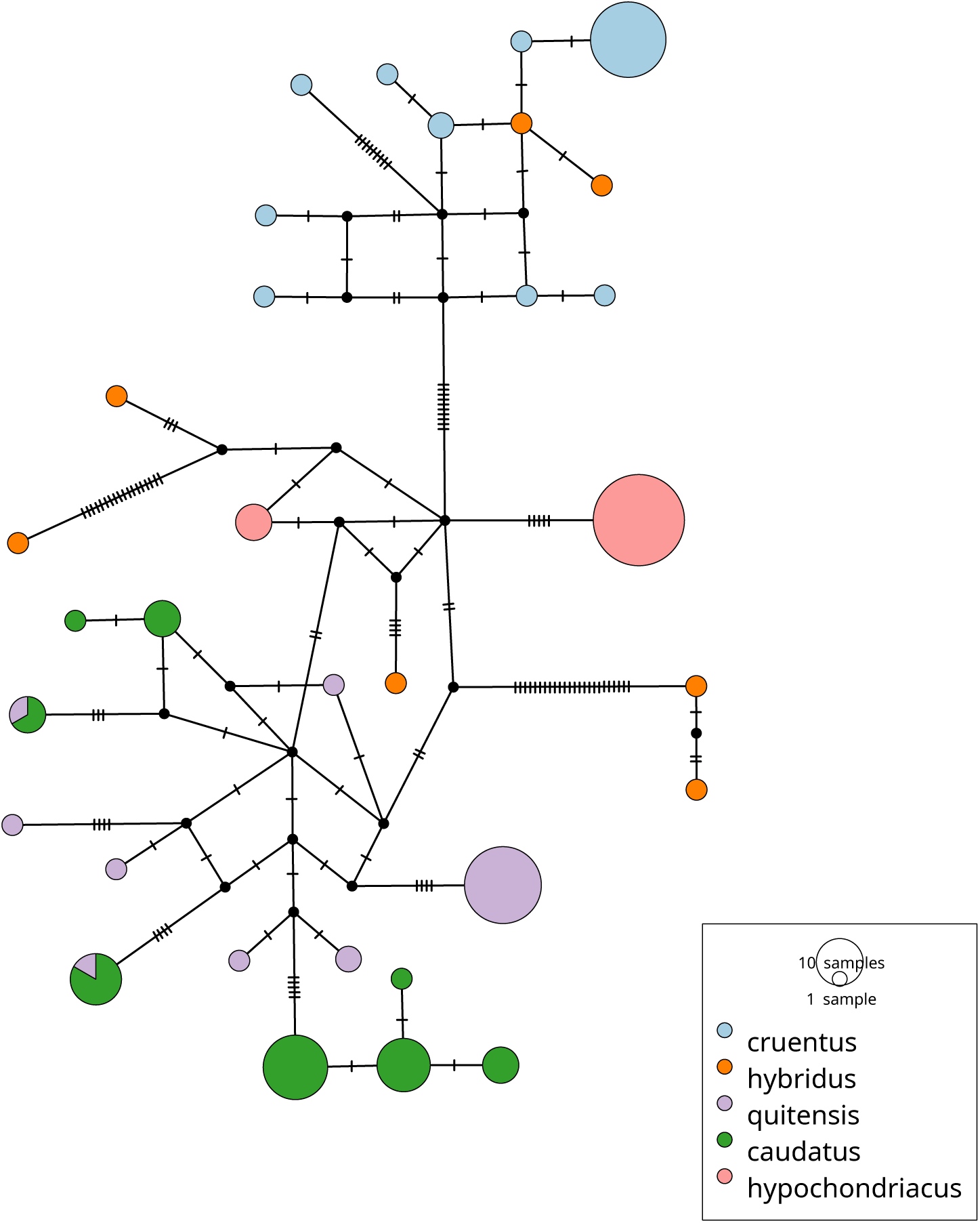
***AmMYBL1* haplotype network from the three domesticates and their wild relatives.** The haplotype network was created from 216 *AmMYBL1* haplotypes from the wild *A. hybridus* and *A. quitensis* and the domesticated *A. hypochondriacus*, *A. caudatus*, and *A. cruentus*. The size of the nodes shows the number of identical haplotypes. Mutations are indicated by marks on the edges. Haplotypes were colored according to species of the accessions.

**Figure S15.**
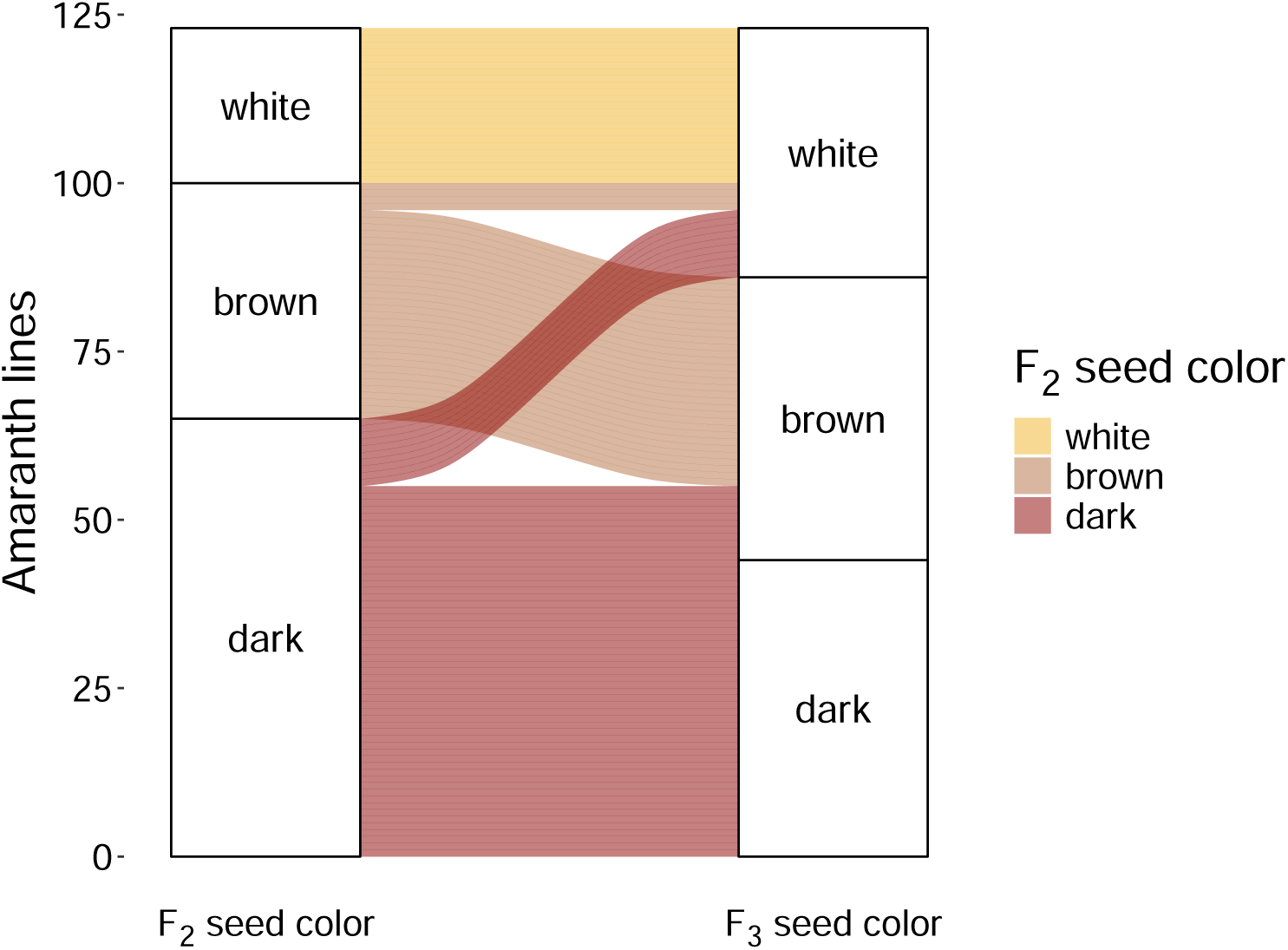
Seed color inheritance in *A. hypochondriacus* F_2_ and F_3_ lines. Changes in seed color from individuals (y-axis) from the F_2_ to F_3_ generation (x-axis) as observed upon self-propagation of the F_2_ lines (e.g., while all F_2_ lines with white seeds produced white seeds also in the F_3_, multiple dark colored F_2_ produced F_3_ offspring lines with brown or white seed color). Colors were chosen to reflect the seed color of the respective F_2_ individual. Note that the propagated F_2_ lines respect a non-random sample from the whole population and their seed color distribution does not reflect the overall distribution of seed colors in the population.

**Figure S16.**
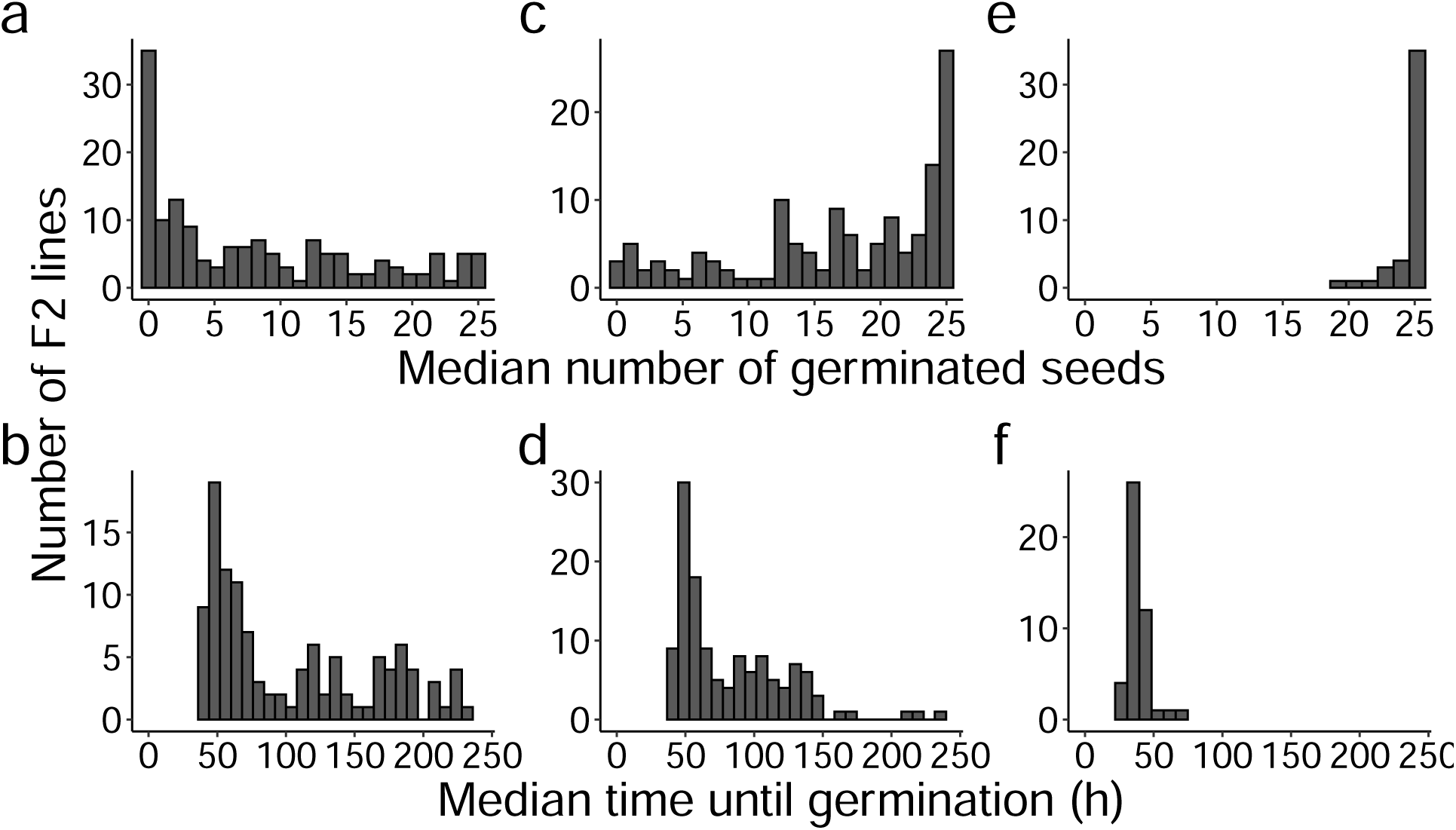
Phenotype distribution in *A. hypochondriacus* germination experiments. Distribution of the median number of germinated seeds and the median time until germination in hours for assessed F_2_ lines of the *A. hypochondriacus* mapping population shortly after harvest (**a**, **b**, n = 150), after four months of storage (**c**, **d**, n = 130), and after 24 months of storage (**e**, **f**, n = 45).

**Figure S17.**
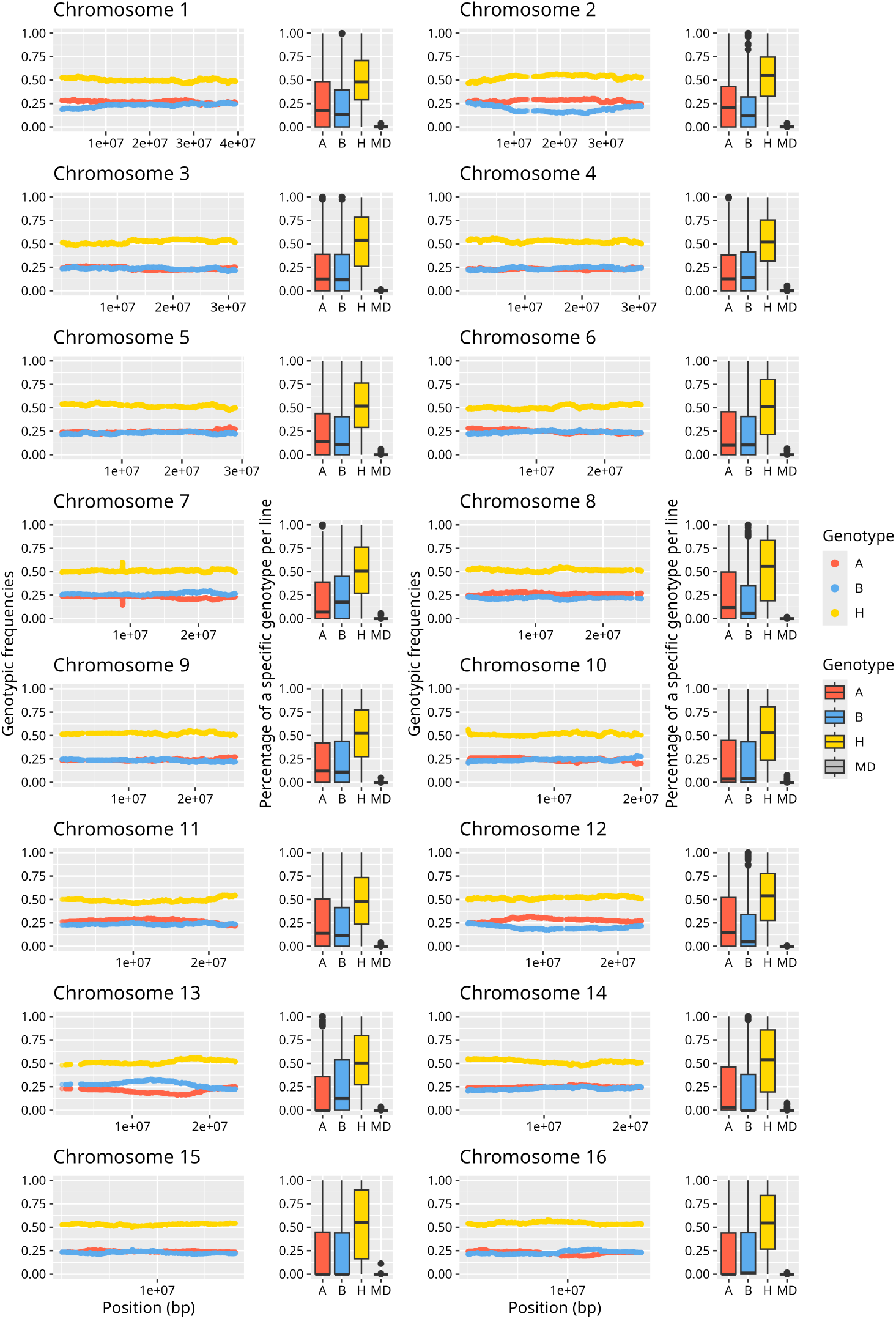
Summary of genotypic frequencies per chromosome of F_2_ lines used for QTL mapping after imputation. For each chromosome, the left side depicts the genotypic frequencies of the homozygous parental (A and B) or heterozygous (H) genotypes per SNP along the chromosome. The right side shows boxplots with per line frequencies of the different genotype calls and missing data (MD) for the respective chromosome.

**Figure S18.**
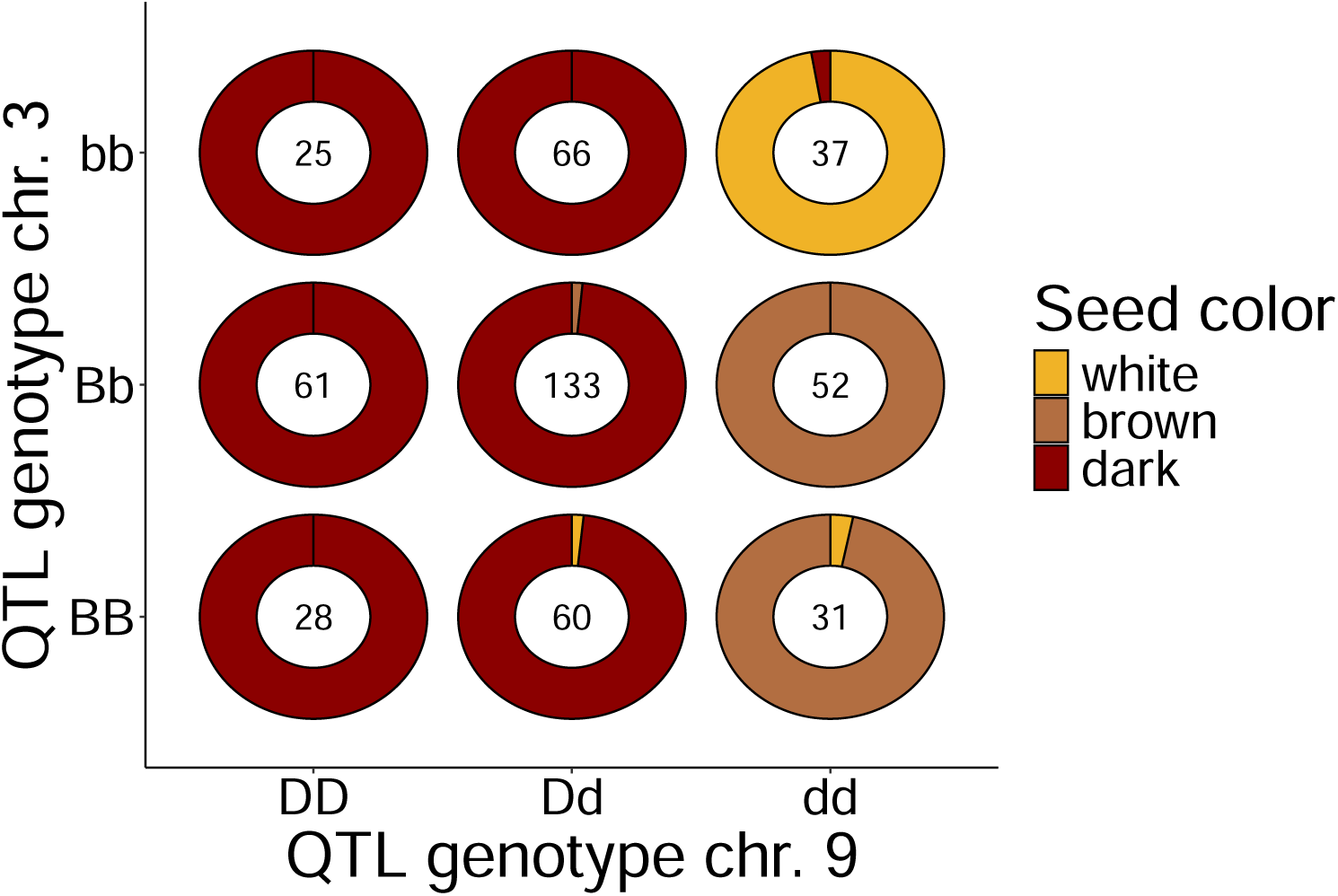
Interaction between seed color phenotype and genotype at QTL peaks in *A. hypochondriacus* F_2_ lines. The genotypes at the chromosome 9 QTL include the dominant dark allele D and the recessive allele d. The genotypes at the chromosome 3 QTL include the hypostatic brown allele B and the recessive allele b. The pie-charts correspond to the fraction of F_2_ individuals with different seed colors, with the central number showing the number of individuals with that combination of genotypes.

**Figure S19.**
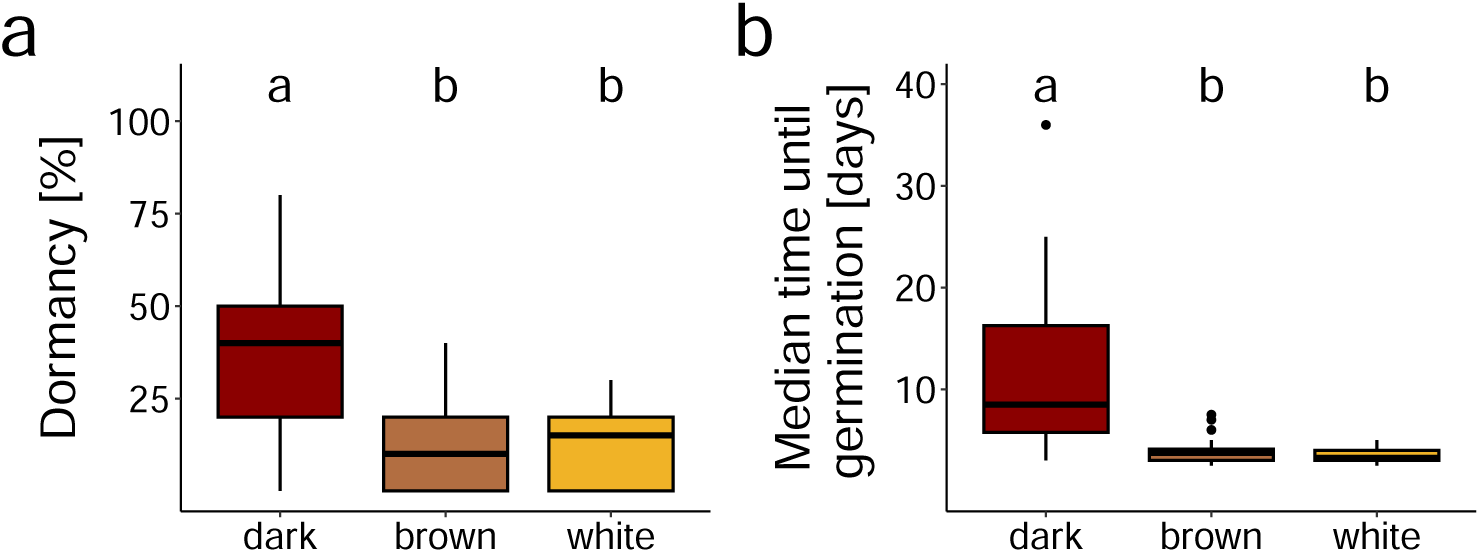
Germination properties of *A. hypochondriacus* F_3_ families in the competition experiment. **a**: Percent of dormant seeds for F_3_ families with different seed colors. Dormant seeds refer to seeds which did not germinate during the experimental duration of the competition experiment. **b**: Median time until germination in days for germinated seeds of F_3_ families with different seed colors. Significant differences between seed colors (p-value < 0.05) between groups are depicted using one-letter representation.

**Figure S20.**
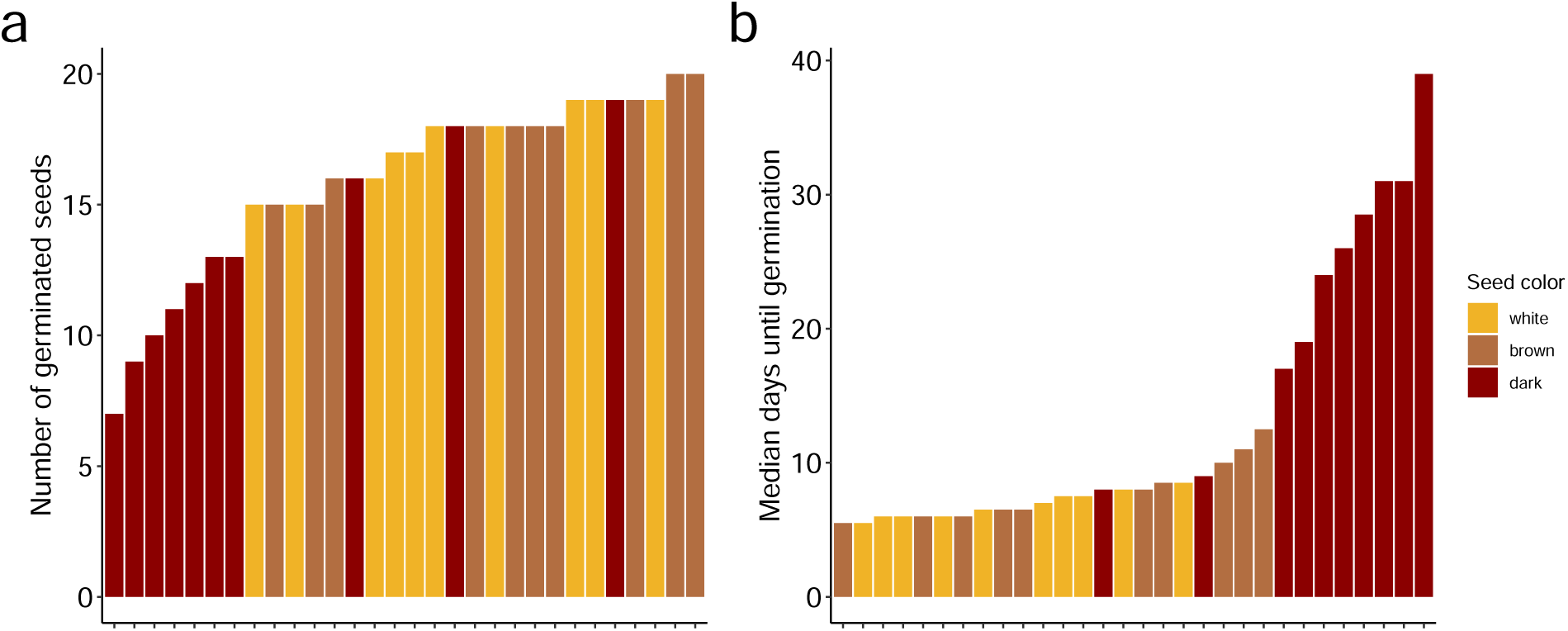
Summary of seed germination metrics during the competition experiment for F_3_ families. **a**: Number of seeds that germinated during the experimental duration per F_3_ family in the experiment. The total number of seeds sown per F_3_ family was 20. **b**: Median days until germination for germinated seeds of F_3_ families during the competition experiment.

**Figure S21.**
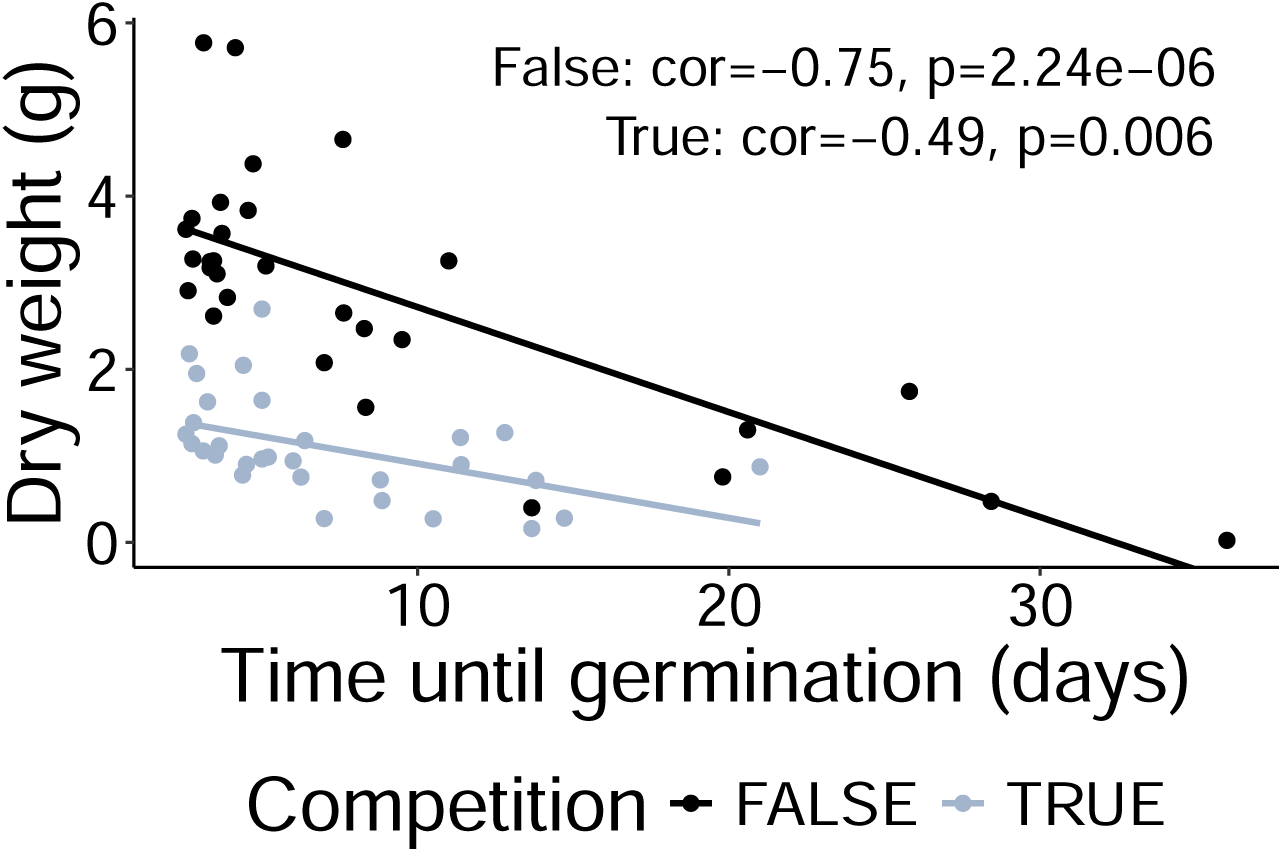
Correlation between germination time and dry weight after 7 weeks for *A. hypochondriacus* F_3_ families in the competition experiment. Pearson’s correlation coefficient between mean dry weight and mean time until germination of F_3_ families (n = 30) grown with or without competition.

**Figure S22.**
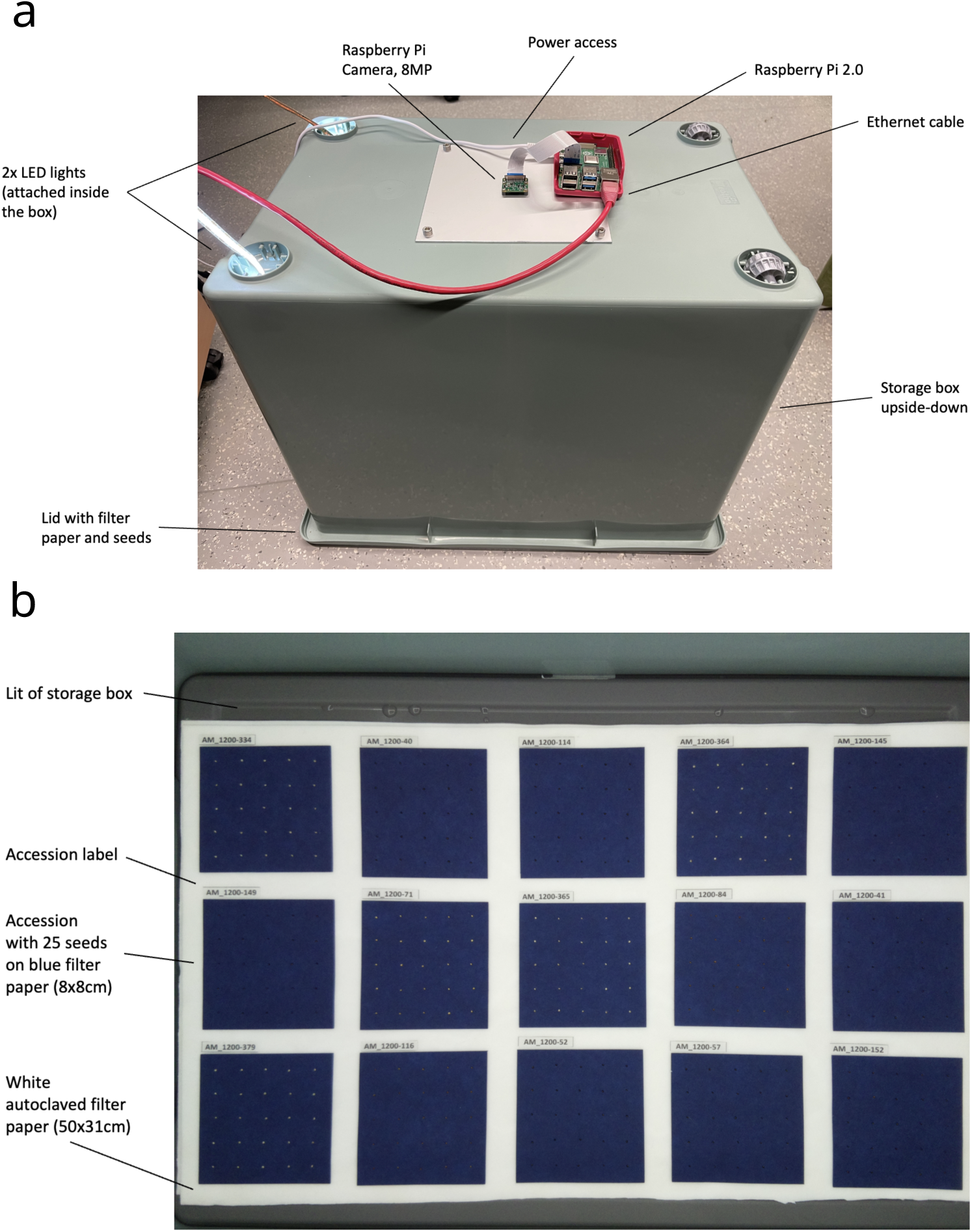
Experimental setup for amaranth germination experiments. Semi-automated setup for the assessment of germination properties of amaranth seeds. **a**: Germination experiments were conducted inside storage boxes to limit water evaporation from wet filter paper with seeds with attached raspberry pi computers controlling the automated imaging every 2 h. Raspberry Pi cameras were fixed to the top of the storage boxes. To enable imaging, constant light was provided by two LED strips attached inside the box. **b**: Image of amaranth seeds arranged in the experimental setup from inside the box taken using the attached Raspberry Pi camera.

**Figure S23.**
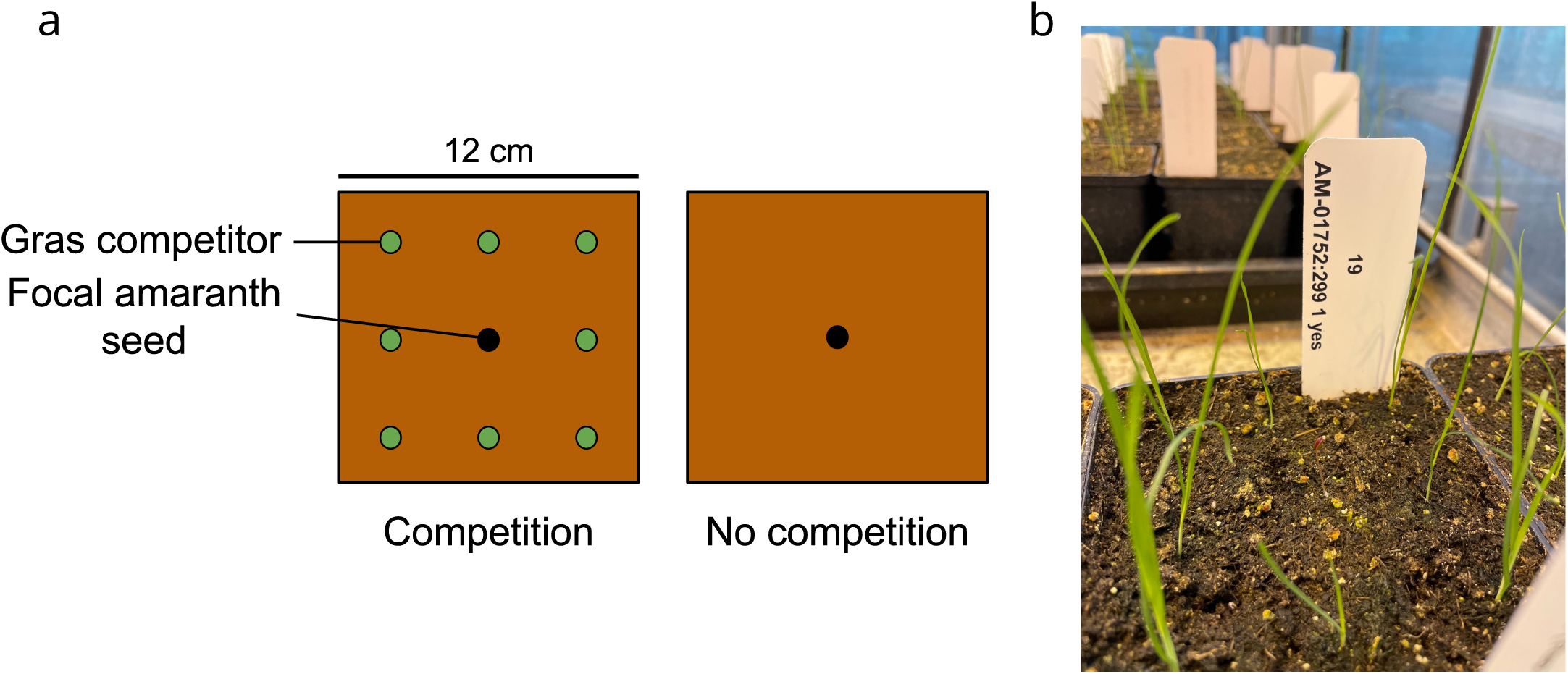
Experimental setup for the *A. hypochondriacus* competition experiment. **a**: Depiction of plant pots used for the experiment. In the competition condition on the left, seedlings of the gras competitor where evenly transplanted around the focal amaranth seed. Without competition, only the amaranth seed was placed in the pot. **b**: Exemplary photo of a germinated focal amaranth seedling surrounded by gras seedlings in the competition condition.

**Figure S24.**
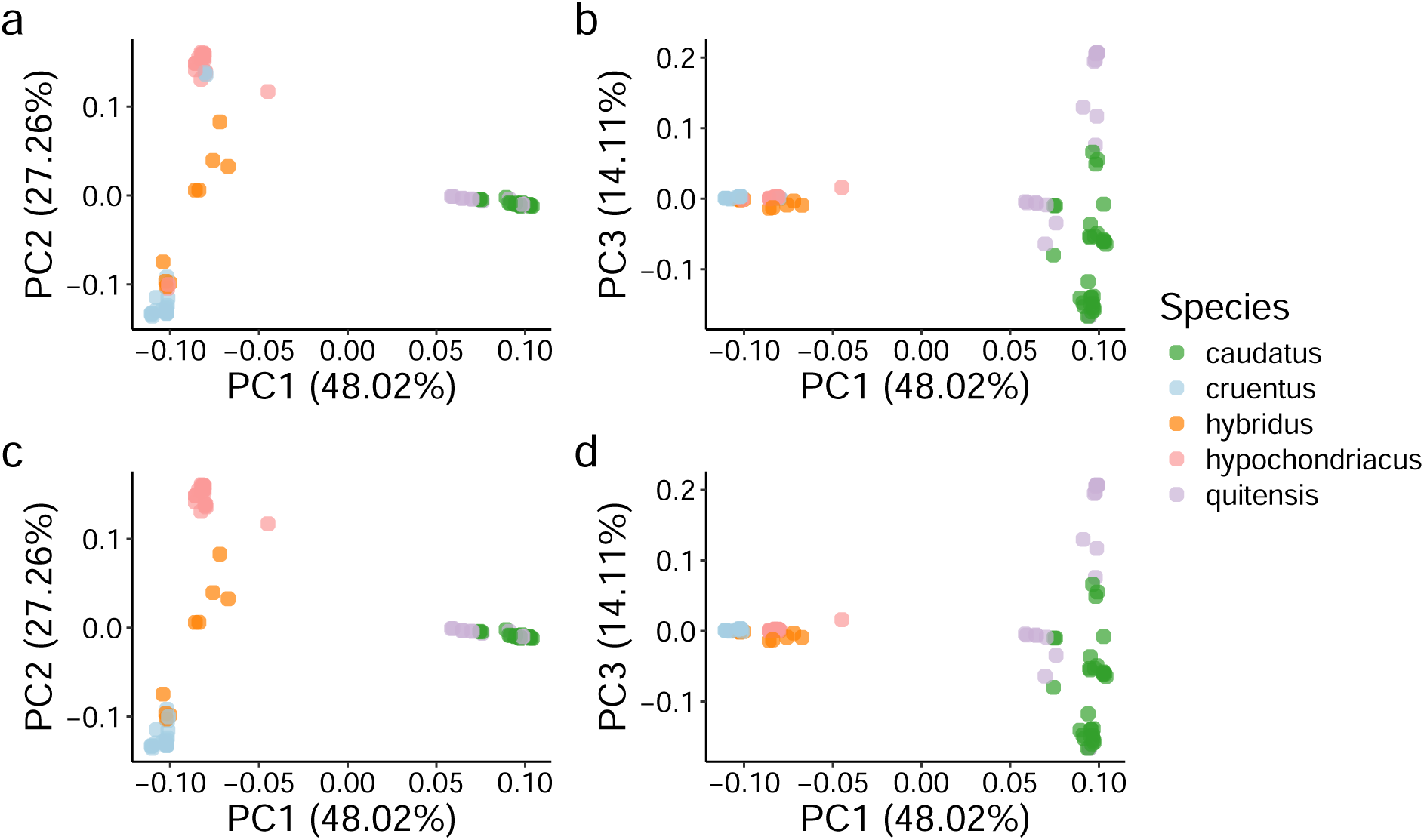
Principal component analysis of whole-genome sequencing data of 115 reclassified *Amaranthus* accessions. Domesticated *Amaranthus* accessions were assigned species and colored according to their clustering in PC1 and PC2. Axis titles include the percent variance explained by the PC. **a**: Clustering of *Amaranthus* accessions on PC1 and PC2 before reclassification. **b**: Clustering of *Amaranthus* accessions on PC1 and PC3 before reclassification. **c**: Clustering of *Amaranthus* accessions on PC1 and PC2 after reclassification. **d**: Clustering of *Amaranthus* accessions on PC1 and PC3 after reclassification.

### Supplementary Tables

**Table S1.**
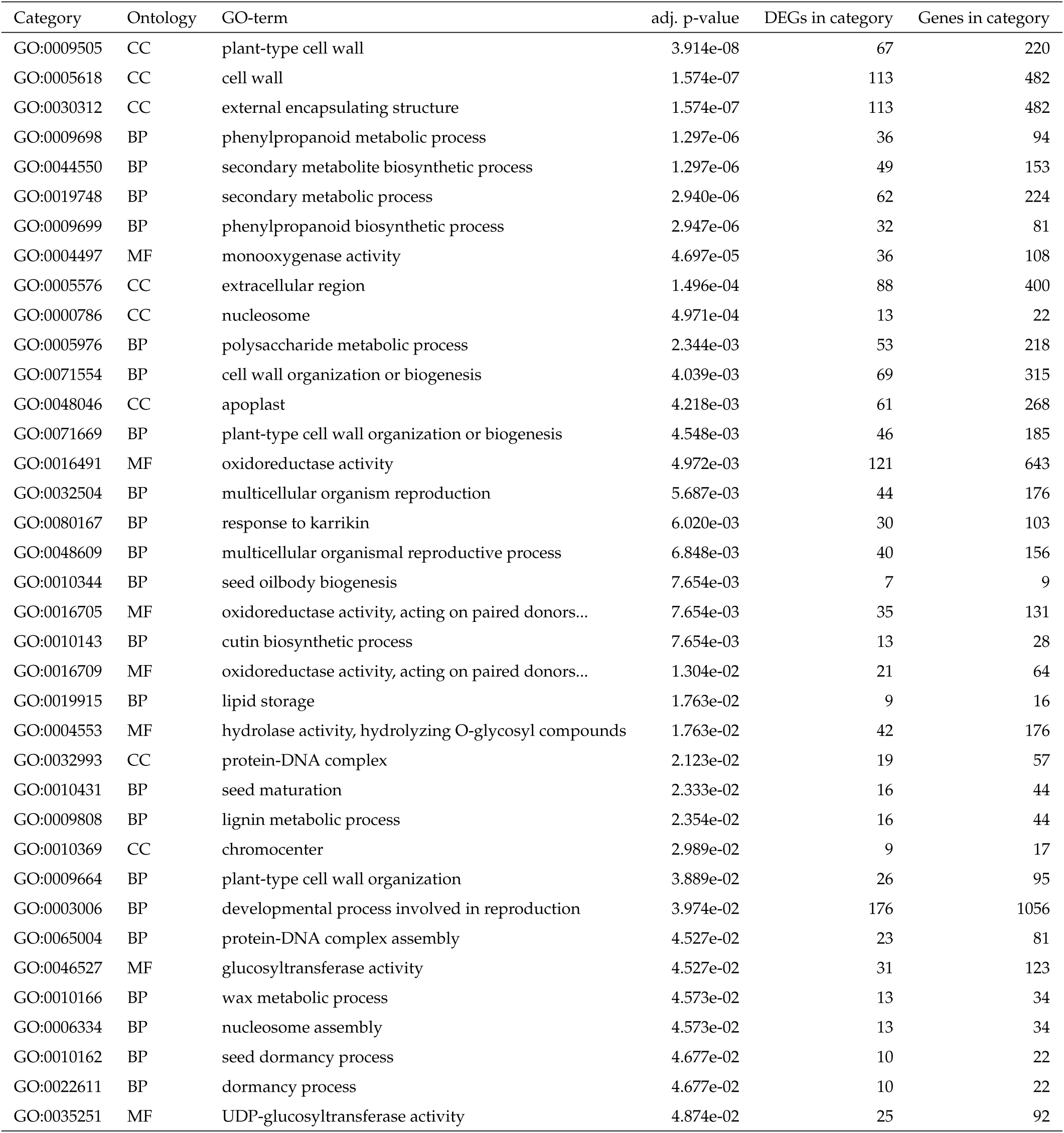
GO-term categories enriched among differentially expressed genes between dark and white developing *A. hypochondriacus* seeds. . For each category, the corresponding ontology, a short description of the GO-term, adjusted p-value for over-representation as well as the number of differentially expressed genes and total genes in the category are displayed.

| Category | Ontology | GO-term | adj. p-value | DEGs in category | Genes in category |
| --- | --- | --- | --- | --- | --- |
| GO:0009505 | CC | plant-type cell wall | 3.914e-08 | 67 | 220 |
| GO:0005618 | CC | cell wall | 1.574e-07 | 113 | 482 |
| GO:0030312 | CC | external encapsulating structure | 1.574e-07 | 113 | 482 |
| GO:0009698 | BP | phenylpropanoid metabolic process | 1.297e-06 | 36 | 94 |
| GO:0044550 | BP | secondary metabolite biosynthetic process | 1.297e-06 | 49 | 153 |
| GO:0019748 | BP | secondary metabolic process | 2.940e-06 | 62 | 224 |
| GO:0009699 | BP | phenylpropanoid biosynthetic process | 2.947e-06 | 32 | 81 |
| GO:0004497 | MF | monooxygenase activity | 4.697e-05 | 36 | 108 |
| GO:0005576 | CC | extracellular region | 1.496e-04 | 88 | 400 |
| GO:0000786 | CC | nucleosome | 4.971e-04 | 13 | 22 |
| GO:0005976 | BP | polysaccharide metabolic process | 2.344e-03 | 53 | 218 |
| GO:0071554 | BP | cell wall organization or biogenesis | 4.039e-03 | 69 | 315 |
| GO:0048046 | CC | apoplast | 4.218e-03 | 61 | 268 |
| GO:0071669 | BP | plant-type cell wall organization or biogenesis | 4.548e-03 | 46 | 185 |
| GO:0016491 | MF | oxidoreductase activity | 4.972e-03 | 121 | 643 |
| GO:0032504 | BP | multicellular organism reproduction | 5.687e-03 | 44 | 176 |
| GO:0080167 | BP | response to karrikin | 6.020e-03 | 30 | 103 |
| GO:0048609 | BP | multicellular organismal reproductive process | 6.848e-03 | 40 | 156 |
| GO:0010344 | BP | seed oilbody biogenesis | 7.654e-03 | 7 | 9 |
| GO:0016705 | MF | oxidoreductase activity, acting on paired donors... | 7.654e-03 | 35 | 131 |
| GO:0010143 | BP | cutin biosynthetic process | 7.654e-03 | 13 | 28 |
| GO:0016709 | MF | oxidoreductase activity, acting on paired donors... | 1.304e-02 | 21 | 64 |
| GO:0019915 | BP | lipid storage | 1.763e-02 | 9 | 16 |
| GO:0004553 | MF | hydrolase activity, hydrolyzing O-glycosyl compounds | 1.763e-02 | 42 | 176 |
| GO:0032993 | CC | protein-DNA complex | 2.123e-02 | 19 | 57 |
| GO:0010431 | BP | seed maturation | 2.333e-02 | 16 | 44 |
| GO:0009808 | BP | lignin metabolic process | 2.354e-02 | 16 | 44 |
| GO:0010369 | CC | chromocenter | 2.989e-02 | 9 | 17 |
| GO:0009664 | BP | plant-type cell wall organization | 3.889e-02 | 26 | 95 |
| GO:0003006 | BP | developmental process involved in reproduction | 3.974e-02 | 176 | 1056 |
| GO:0065004 | BP | protein-DNA complex assembly | 4.527e-02 | 23 | 81 |
| GO:0046527 | MF | glucosyltransferase activity | 4.527e-02 | 31 | 123 |
| GO:0010166 | BP | wax metabolic process | 4.573e-02 | 13 | 34 |
| GO:0006334 | BP | nucleosome assembly | 4.573e-02 | 13 | 34 |
| GO:0010162 | BP | seed dormancy process | 4.677e-02 | 10 | 22 |
| GO:0022611 | BP | dormancy process | 4.677e-02 | 10 | 22 |
| GO:0035251 | MF | UDP-glucosyltransferase activity | 4.874e-02 | 25 | 92 |

**Table S2.**
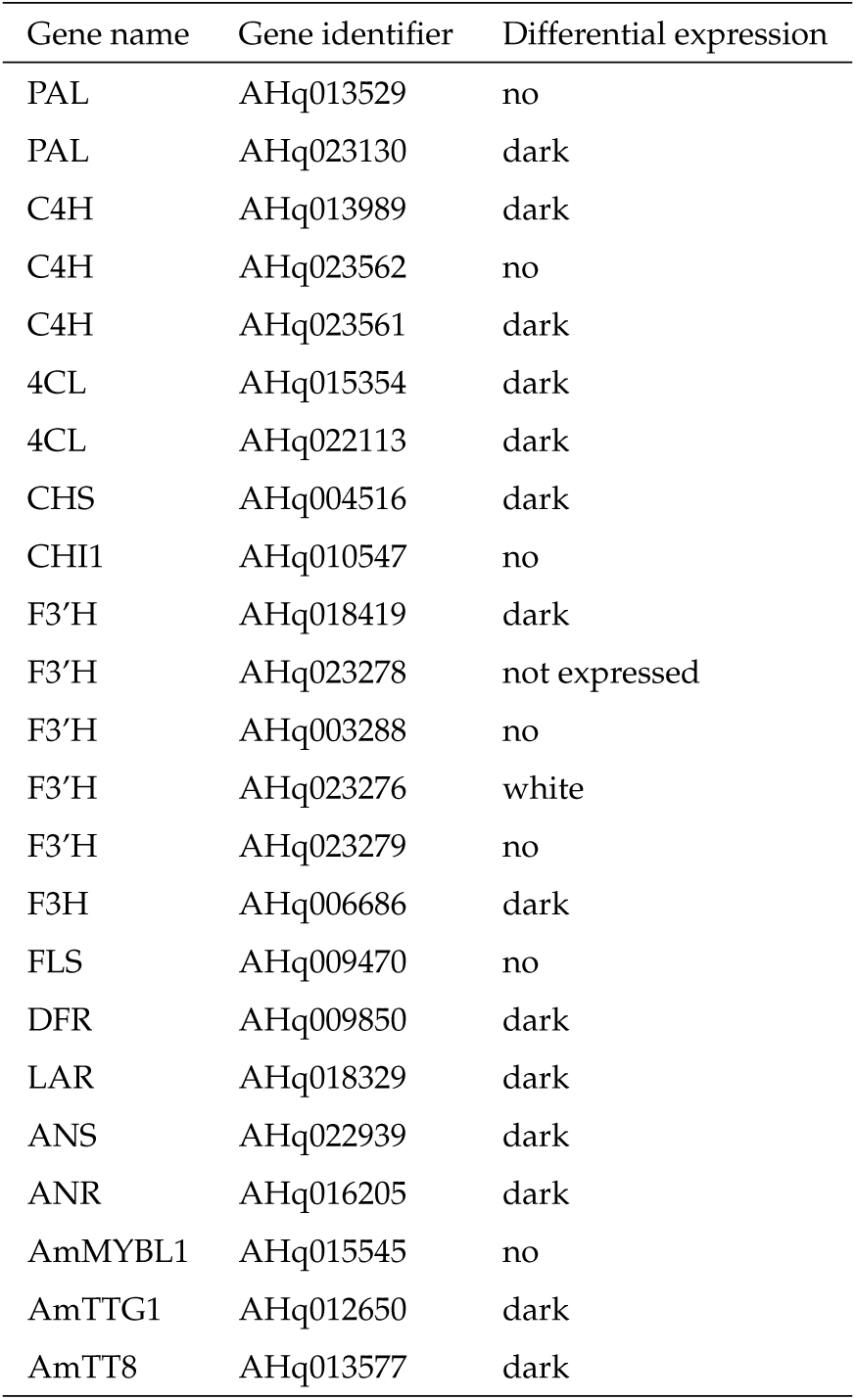
Expression status of flavonoid pathway genes in *A. hypochondriacus* seeds. . The table includes gene name, identifier, and differential expression status of amaranth flavonoid pathway genes in dark and white seeds. Significantly increased expression in the two seed colors is annotated as “dark” or “white” for increased expression in dark or white seeds, respectively, while “no” indicates no difference in gene expression between seed colors. One gene copy of *F3’H* (AHq023278) was not expressed in the seeds.

**Table S3.** PCR-based genotyping of the TE insertion in *AmMYBL1* in *A. hypochondriacus* accessions. TE genotype was ass<u>igned based on 421 bp length polymorphism</u> in PCR.

| Accession | Species | Seed color | TE genotype |
| --- | --- | --- | --- |
| Ames 15204 | *A. hypochondriacus* | dark | no TE |
| Ames 2257 | *A. hypochondriacus* | dark | no TE |
| Ames 5585 | *A. hypochondriacus* | dark | no TE |
| Ames 5586 | *A. hypochondriacus* | dark | no TE |
| Ames 5603 | *A. hypochondriacus* | dark | no TE |
| Ames 5611 | *A. hypochondriacus* | dark | no TE |
| Ames 5612 | *A. hypochondriacus* | dark | no TE |
| PI 274278 | *A. hypochondriacus* | dark | no TE |
| PI 481042 | *A. hypochondriacus* | dark | no TE |
| PI 481075 | *A. hypochondriacus* | dark | no TE |
| PI 481076 | *A. hypochondriacus* | dark | no TE |
| PI 481091 | *A. hypochondriacus* | dark | no TE |
| PI 481107 | *A. hypochondriacus* | dark | no TE |
| PI 481108 | *A. hypochondriacus* | dark | no TE |
| PI 481110 | *A. hypochondriacus* | dark | no TE |
| PI 481184 | *A. hypochondriacus* | dark | no TE |
| PI 481336 | *A. hypochondriacus* | dark | no TE |
| PI 669884 | *A. hypochondriacus* | dark | no TE |
| Ames 15210 | *A. hypochondriacus* | white | TE insertion |
| Ames 15301 | *A. hypochondriacus* | white | TE insertion |
| Ames 5138 | *A. hypochondriacus* | white | TE insertion |
| Ames 5145 | *A. hypochondriacus* | white | TE insertion |
| Ames 5587 | *A. hypochondriacus* | white | TE insertion |
| Ames 5588 | *A. hypochondriacus* | white | TE insertion |
| Ames 5593 | *A. hypochondriacus* | white | TE insertion |
| Ames 5608 | *A. hypochondriacus* | white | TE insertion |
| PI 175040 | *A. hypochondriacus* | white | TE insertion |
| PI 274277 | *A. hypochondriacus* | white | TE insertion |
| PI 274279 | *A. hypochondriacus* | white | TE insertion |
| PI 480557 | *A. hypochondriacus* | white | TE insertion |
| PI 480569 | *A. hypochondriacus* | white | TE insertion |
| PI 481074 | *A. hypochondriacus* | white | TE insertion |
| PI 481093 | *A. hypochondriacus* | white | TE insertion |
| PI 481106 | *A. hypochondriacus* | white | TE insertion |
| PI 481109 | *A. hypochondriacus* | white | TE insertion |
| PI 615696 | *A. hypochondriacus* | white | TE insertion |
| PI 636183 | *A. hypochondriacus* | white | TE insertion |
| PI 636184 | *A. hypochondriacus* | white | TE insertion |
| PI 636185 | *A. hypochondriacus* | white | TE insertion |
| PI 636186 | *A. hypochondriacus* | white | TE insertion |
| PI 636187 | *A. hypochondriacus* | white | TE insertion |
| PI 636188 | *A. hypochondriacus* | white | TE insertion |
| PI 636190 | *A. hypochondriacus* | white | TE insertion |
| PI 636191 | *A. hypochondriacus* | white | TE insertion |
| PI 636192 | *A. hypochondriacus* | white | TE insertion |
| PI 669931 | *A. hypochondriacus* | white | TE insertion |
| PI 674256 | *A. hypochondriacus* | white | TE insertion |
| PI 689735 | *A. hypochondriacus* | white | TE insertion |
| PI 689736 | *A. hypochondriacus* | white | TE insertion |
| PI 689738 | *A. hypochondriacus* | white | TE insertion |

**Table S4.** Summary of the seed color of interspecific *Amaranthus* crosses. Detailed are the accession identifiers, species and seed color of maternal and paternal accessions and the seed color the <u>offspring F_1_ plants produced.</u>

| Maternal accession | Maternal species | Maternal seed color | Paternal accession | Paternal species | Paternal seed color | Offspring seed color |
| --- | --- | --- | --- | --- | --- | --- |
| PI 490518 | *A. caudatus* | dark | PI 558499 | *A. hypochondriacus* | white | dark |
| PI 490518 | *A. caudatus* | dark | PI 643070 | *A. hypochondriacus* | white | dark |
| PI 490612 | *A. caudatus* | white | PI 643070 | *A. hypochondriacus* | white | white |
| PI 490612 | *A. caudatus* | white | PI 558499 | *A. hypochondriacus* | white | white |
| PI 558499 | *A. hypochondriacus* | white | PI 643058 | *A. cruentus* | white | white |

**Table S5.** Chi-square tests for different inheritance models for seed color segregation patterns in the *A. hypochondriacus* F_2_ mapping population. Seed colors and the observed segregation ratio for 511 F_2_ lines are depicted, alongside chi-square tests to test whether the observed ratios fit the expected distributions for two-locus inheritance models. The observed segregation ratio fit neither to recessive epistasis nor to duplicate interaction inheritance models, whereas segregation according to dominant epistasis between the two loci could not be rejected.

|  | Observed seed colors | Observed ratios | Dominant epistasis | Recessive epistasis | Duplicate interaction |
| --- | --- | --- | --- | --- | --- |
| Dark | 385 | 12.05 | 12 | 9 | 9 |
| Brown | 87 | 2.72 | 3 | 4 | 6 |
| White | 39 | 1.22 | 1 | 3 | 1 |
| Chisq. p-value |  |  | 0.304 | 0 | 0 |

**Table S6.** Summary of *Amaranthus* samples used for metabolomic analysis. Detailed are the sample identifier, accession name, species and seed color of accessions used for untargeted metabolomic analysis.

| Sample | Accession | Species | Seed color |
| --- | --- | --- | --- |
| AM_01103 | PI 643070 | *A. hypochondriacus* | white |
| AM_02313 | PI 643070 | *A. hypochondriacus* | white |
| AM_02259 | PI 643070 | *A. hypochondriacus* | white |
| AM_02261 | PI 558499 | *A. hypochondriacus* | white |
| AM_02262 | PI 558499 | *A. hypochondriacus* | white |
| AM_02263 | PI 558499 | *A. hypochondriacus* | white |
| AM_01723 | PI 604587 | *A. hypochondriacus* | dark |
| AM_02283 | PI 604587 | *A. hypochondriacus* | dark |
| AM_02312 | PI 604587 | *A. hypochondriacus* | dark |
| AM_02314 | PI 604581 | *A. hypochondriacus* | dark |
| AM_02315 | PI 604581 | *A. hypochondriacus* | dark |
| AM_02316 | PI 604581 | *A. hypochondriacus* | dark |
| AM_02292 | PI 511754 | *A. hybridus* | dark |
| AM_02293 | PI 511754 | *A. hybridus* | dark |
| AM_02294 | PI 511754 | *A. hybridus* | dark |
| AM_02298 | PI 667158 | *A. hybridus* | dark |
| AM_02299 | PI 667158 | *A. hybridus* | dark |
| AM_02300 | PI 667158 | *A. hybridus* | dark |

**Table S7.**
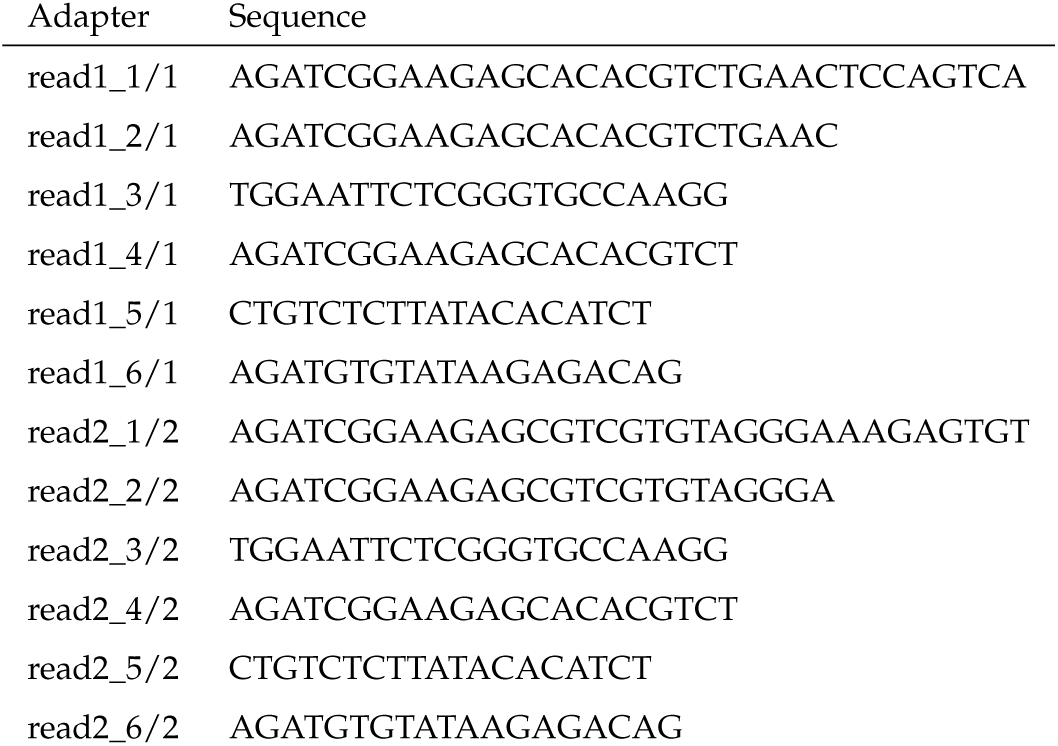
Adapter sequences used for adapter trimming of RNA sequencing reads generated from developing *A. hypochondriacus* seeds.

**Table S8.**
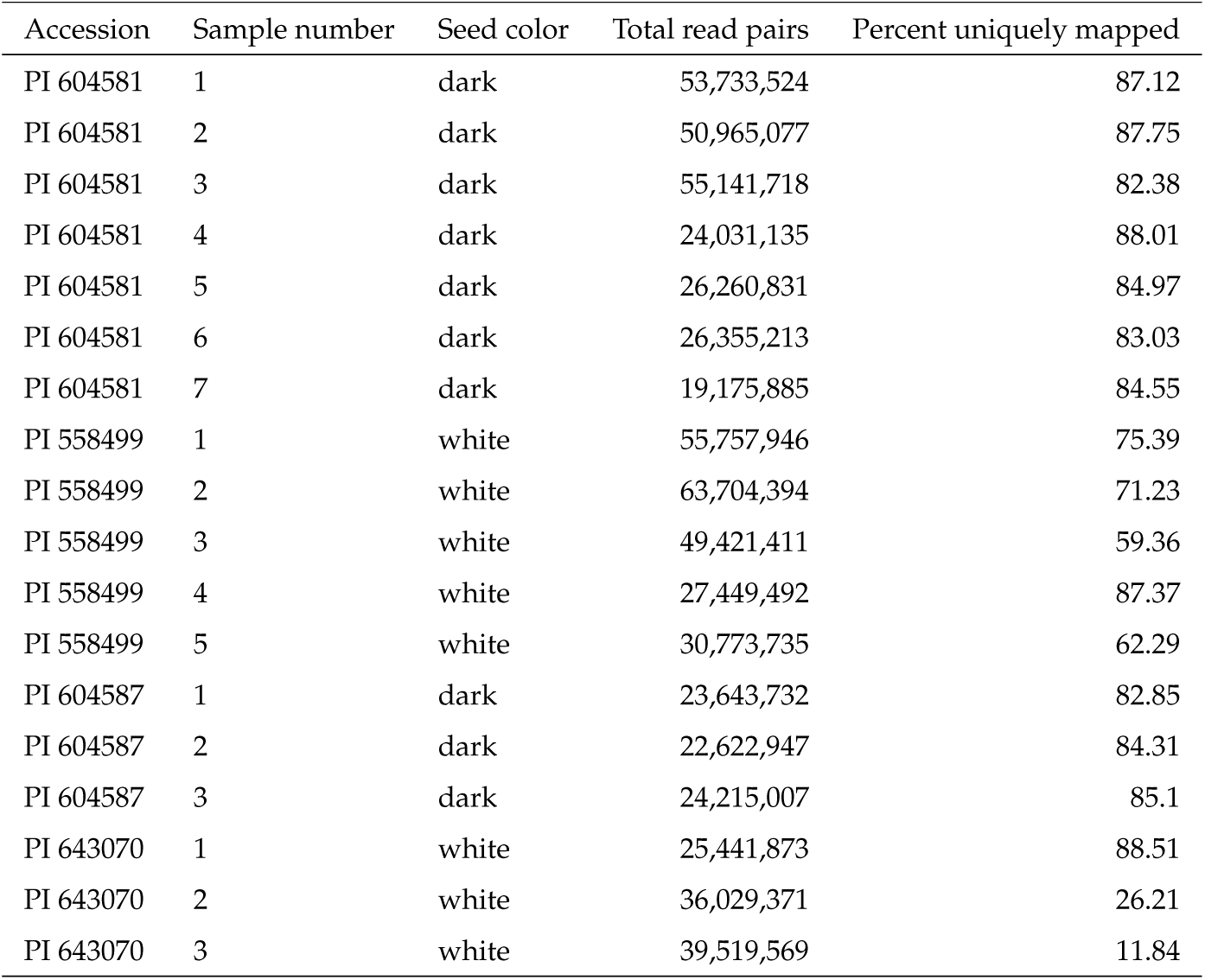
Sequencing statistics for samples used for differential expression analysis. Displayed are the accession, seed color, number of total read pairs, and the percentage of uniquely mapped reads for each sample used for the differential expression analysis.

**Table S9.**
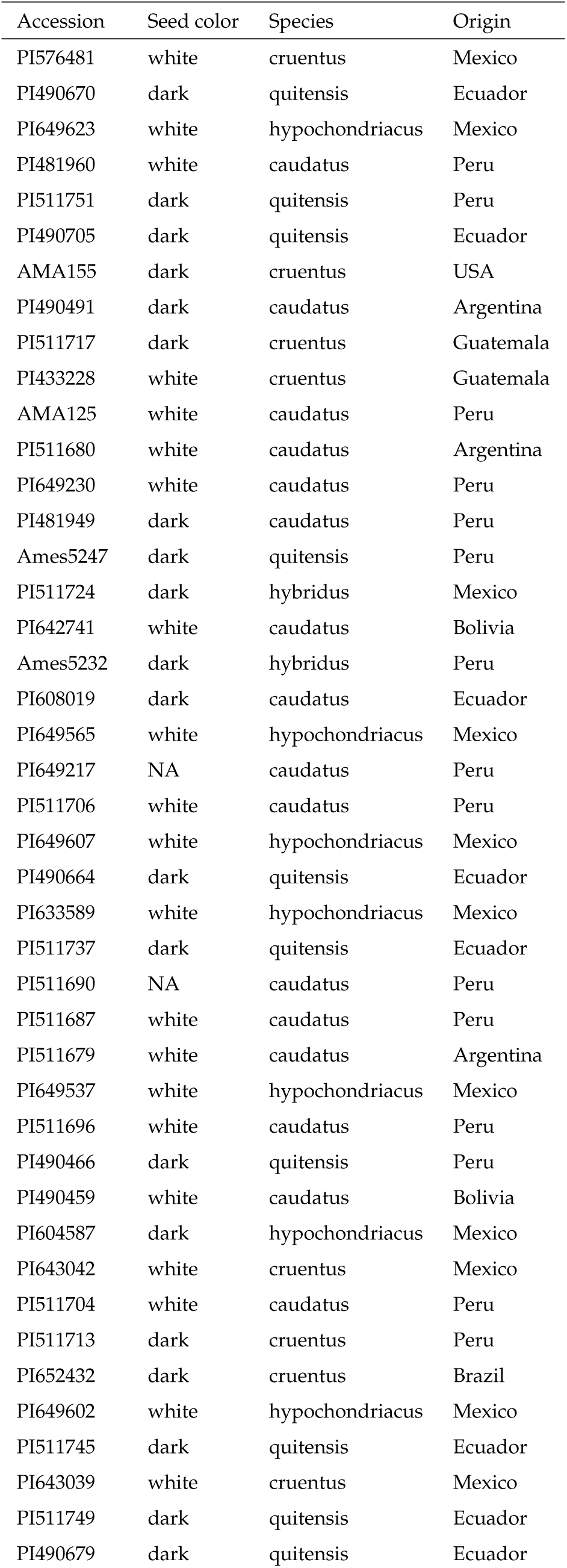
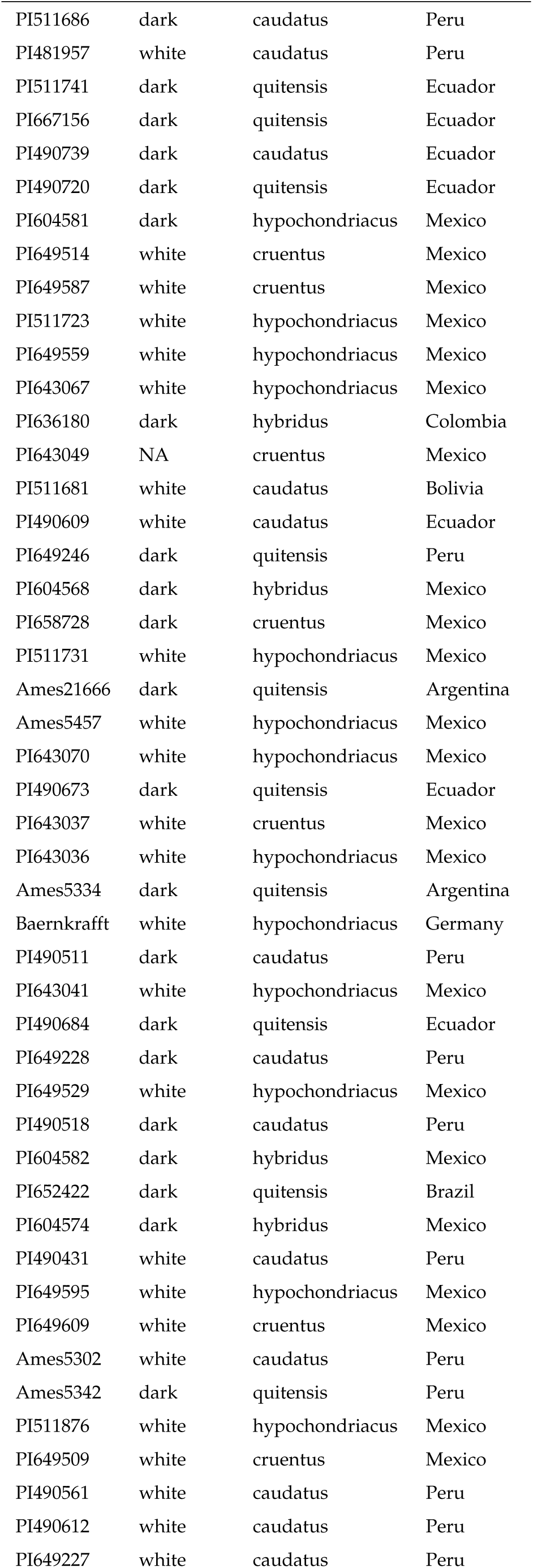
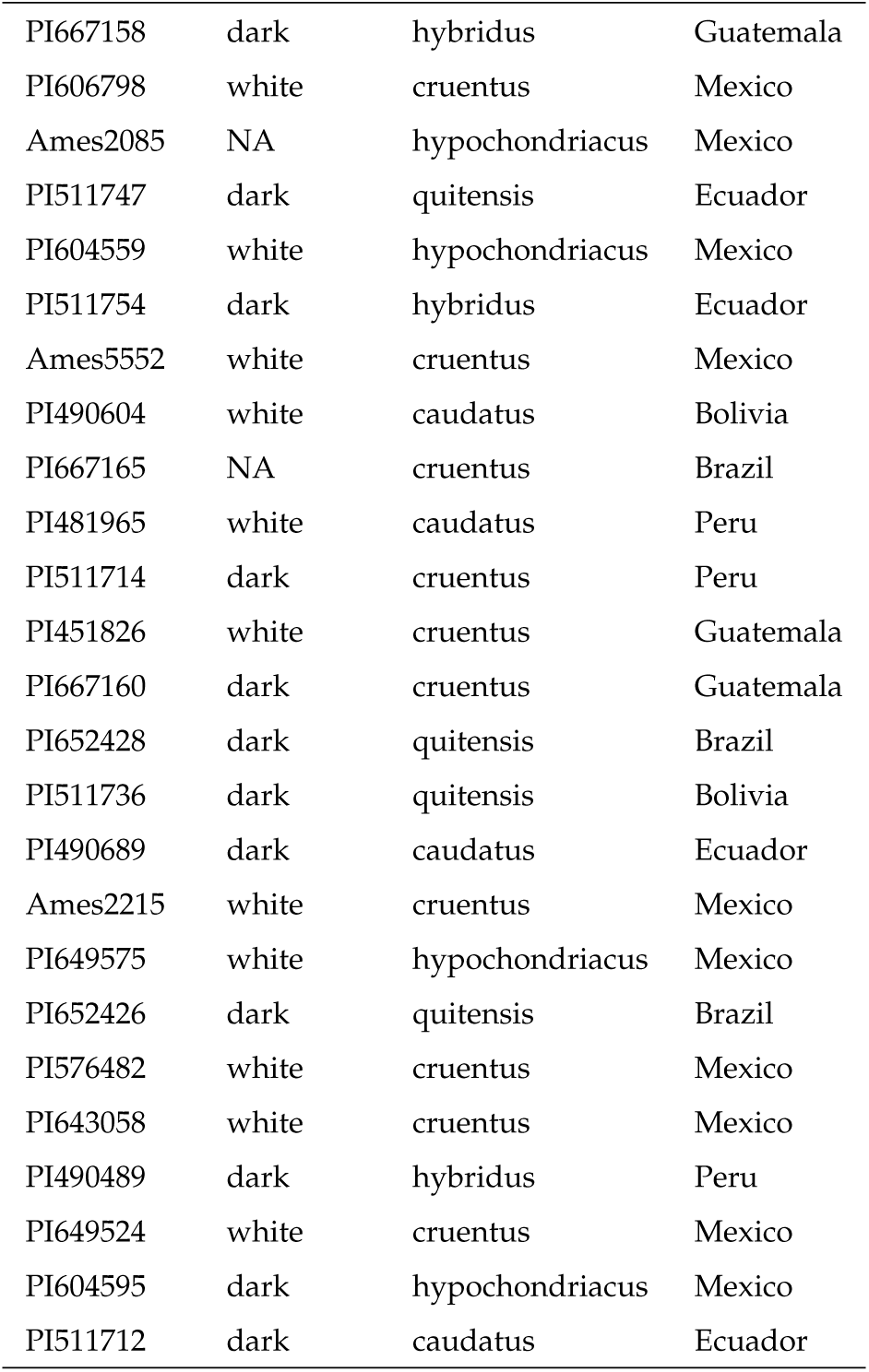
List of reclassified grain amaranth accessions used for genomic analysis. We reassigned the species identifier based on clustering of the accessions in the PCA of genome-wide bi-allelic SNPs.

| Accession | Seed color | Species | Origin |
| --- | --- | --- | --- |
| PI576481 | white | cruentus | Mexico |
| PI490670 | dark | quitensis | Ecuador |
| PI649623 | white | hypochondriacus | Mexico |
| PI481960 | white | caudatus | Peru |
| PI511751 | dark | quitensis | Peru |
| PI490705 | dark | quitensis | Ecuador |
| AMA155 | dark | cruentus | USA |
| PI490491 | dark | caudatus | Argentina |
| PI511717 | dark | cruentus | Guatemala |
| PI433228 | white | cruentus | Guatemala |
| AMA125 | white | caudatus | Peru |
| PI511680 | white | caudatus | Argentina |
| PI649230 | white | caudatus | Peru |
| PI481949 | dark | caudatus | Peru |
| Ames5247 | dark | quitensis | Peru |
| PI511724 | dark | hybridus | Mexico |
| PI642741 | white | caudatus | Bolivia |
| Ames5232 | dark | hybridus | Peru |
| PI608019 | dark | caudatus | Ecuador |
| PI649565 | white | hypochondriacus | Mexico |
| PI649217 | NA | caudatus | Peru |
| PI511706 | white | caudatus | Peru |
| PI649607 | white | hypochondriacus | Mexico |
| PI490664 | dark | quitensis | Ecuador |
| PI633589 | white | hypochondriacus | Mexico |
| PI511737 | dark | quitensis | Ecuador |
| PI511690 | NA | caudatus | Peru |
| PI511687 | white | caudatus | Peru |
| PI511679 | white | caudatus | Argentina |
| PI649537 | white | hypochondriacus | Mexico |
| PI511696 | white | caudatus | Peru |
| PI490466 | dark | quitensis | Peru |
| PI490459 | white | caudatus | Bolivia |
| PI604587 | dark | hypochondriacus | Mexico |
| PI643042 | white | cruentus | Mexico |
| PI511704 | white | caudatus | Peru |
| PI511713 | dark | cruentus | Peru |
| PI652432 | dark | cruentus | Brazil |
| PI649602 | white | hypochondriacus | Mexico |
| PI511745 | dark | quitensis | Ecuador |
| PI643039 | white | cruentus | Mexico |
| PI511749 | dark | quitensis | Ecuador |
| PI490679 | dark | quitensis | Ecuador |
| PI511686 | dark | caudatus | Peru |
| PI481957 | white | caudatus | Peru |
| PI511741 | dark | quitensis | Ecuador |
| PI667156 | dark | quitensis | Ecuador |
| PI490739 | dark | caudatus | Ecuador |
| PI490720 | dark | quitensis | Ecuador |
| PI604581 | dark | hypochondriacus | Mexico |
| PI649514 | white | cruentus | Mexico |
| PI649587 | white | cruentus | Mexico |
| PI511723 | white | hypochondriacus | Mexico |
| PI649559 | white | hypochondriacus | Mexico |
| PI643067 | white | hypochondriacus | Mexico |
| PI636180 | dark | hybridus | Colombia |
| PI643049 | NA | cruentus | Mexico |
| PI511681 | white | caudatus | Bolivia |
| PI490609 | white | caudatus | Ecuador |
| PI649246 | dark | quitensis | Peru |
| PI604568 | dark | hybridus | Mexico |
| PI658728 | dark | cruentus | Mexico |
| PI511731 | white | hypochondriacus | Mexico |
| Ames21666 | dark | quitensis | Argentina |
| Ames5457 | white | hypochondriacus | Mexico |
| PI643070 | white | hypochondriacus | Mexico |
| PI490673 | dark | quitensis | Ecuador |
| PI643037 | white | cruentus | Mexico |
| PI643036 | white | hypochondriacus | Mexico |
| Ames5334 | dark | quitensis | Argentina |
| Baernkrafft | white | hypochondriacus | Germany |
| PI490511 | dark | caudatus | Peru |
| PI643041 | white | hypochondriacus | Mexico |
| PI490684 | dark | quitensis | Ecuador |
| PI649228 | dark | caudatus | Peru |
| PI649529 | white | hypochondriacus | Mexico |
| PI490518 | dark | caudatus | Peru |
| PI604582 | dark | hybridus | Mexico |
| PI652422 | dark | quitensis | Brazil |
| PI604574 | dark | hybridus | Mexico |
| PI490431 | white | caudatus | Peru |
| PI649595 | white | hypochondriacus | Mexico |
| PI649609 | white | cruentus | Mexico |
| Ames5302 | white | caudatus | Peru |
| Ames5342 | dark | quitensis | Peru |
| PI511876 | white | hypochondriacus | Mexico |
| PI649509 | white | cruentus | Mexico |
| PI490561 | white | caudatus | Peru |
| PI490612 | white | caudatus | Peru |
| PI649227 | white | caudatus | Peru |
| PI667158 | dark | hybridus | Guatemala |
| PI606798 | white | cruentus | Mexico |
| Ames2085 | NA | hypochondriacus | Mexico |
| PI511747 | dark | quitensis | Ecuador |
| PI604559 | white | hypochondriacus | Mexico |
| PI511754 | dark | hybridus | Ecuador |
| Ames5552 | white | cruentus | Mexico |
| PI490604 | white | caudatus | Bolivia |
| PI667165 | NA | cruentus | Brazil |
| PI481965 | white | caudatus | Peru |
| PI511714 | dark | cruentus | Peru |
| PI451826 | white | cruentus | Guatemala |
| PI667160 | dark | cruentus | Guatemala |
| PI652428 | dark | quitensis | Brazil |
| PI511736 | dark | quitensis | Bolivia |
| PI490689 | dark | caudatus | Ecuador |
| Ames2215 | white | cruentus | Mexico |
| PI649575 | white | hypochondriacus | Mexico |
| PI652426 | dark | quitensis | Brazil |
| PI576482 | white | cruentus | Mexico |
| PI643058 | white | cruentus | Mexico |
| PI490489 | dark | hybridus | Peru |
| PI649524 | white | cruentus | Mexico |
| PI604595 | dark | hypochondriacus | Mexico |
| PI511712 | dark | caudatus | Ecuador |

**Table S10.**
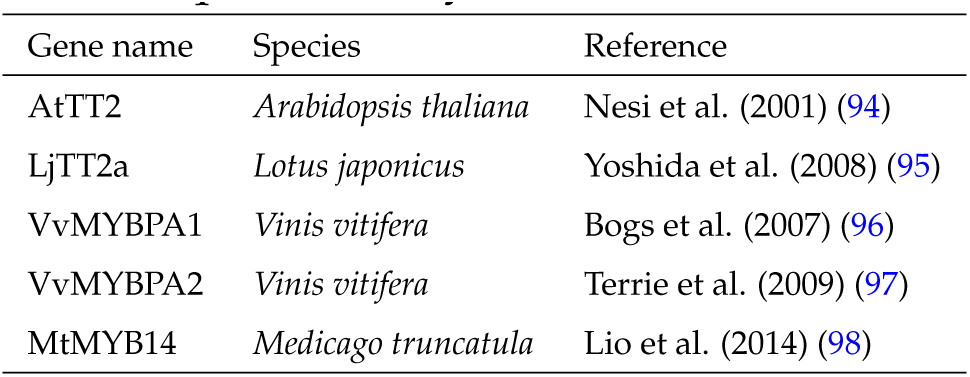
Proanthocyanidin MYB transcription factors from other species used for phylogenetic analysis with *AmMYBL1*. Depicted are gene name, species, and reference publication of proteins included in the phylogenetic and sequence analysis.

